# Self-regulation of neuronal activity in prefrontal cortex

**DOI:** 10.64898/2026.08.18.742749

**Authors:** Chris S. Ki, Ryan C. Williamson, Akash Umakantha, Byron M. Yu, Matthew A. Smith

## Abstract

Despite our best efforts to stay focused on a task, our arousal waxes and wanes over time. Lower levels of arousal are typically associated with drowsiness, whereas higher levels are often associated with stress. These changes in arousal move us away from ideal task performance and manifest as fluctuations in neural activity. We asked whether moment-by-moment neurofeedback could be used to counteract neural fluctuations and thereby regulate arousal levels. Here, we developed an intracortical brain-computer interface (BCI) in which animals used visual neurofeedback to maintain neural population activity in prefrontal cortex near a pre-specified activity target. We found animals used moment-to-moment neurofeedback to reduce neural fluctuations on timescales of seconds to hundreds of milliseconds, and that arousal-related regulation of neural activity was associated with BCI use. Our findings suggest that neurofeedback may enhance or restore regulation of neural activity, with potential clinical applications in conditions where such regulation is impaired.

## Introduction

Even when presented with the same stimulus multiple times, our behavioral responses vary. This behavioral variability may arise in part from fluctuations in internal states. Internal states capture aspects of brain function that cannot be observed directly, but can be inferred from behavior, physiological signals, and changes in neural activity (Flavell *et al.* [1]). This approach has been used to describe internal states such as attention (Cohen *et al.* [2]), decision-related variables (Kiani *et al.* [3]), arousal (McGinley *et al.* [4], Cowley *et al.* [5], Hennig *et al.* [6], Stringer *et al.* [7]), motivation (Allen *et al.* [8]), anxiety (Kennedy *et al.* [9]), and aggression (Nair *et al.* [10]). As these states fluctuate over time, they can influence how sensory inputs are processed and how behavioral responses are generated.

For example, consider a student attending a lecture. At the start of class, their arousal level may be high, helping them stay focused. This state is reflected in their large pupil diameter and corresponds to initial population activity (Fig. 1A, green dot) that lies within a region of neural space representing high arousal levels (Fig. 1A, green sphere). As the lecture progresses, the student’s arousal level may decline, making it harder to stay focused. This decline is reflected by a smaller pupil diameter and a drift in neural activity away from the initial population activity and out of this high arousal region (Fig. 1B, black dashed arrow). If it were possible to volitionally maintain neural activity within or near this target region (Fig. 1C, black dashed arrow), the student might be better able to maintain focus. Many tasks presumably have analogous target activity regions, corresponding to a set of internal states well-suited for task performance. Because variations in internal states contribute to behavioral variability, maintaining specific internal states, such as heightened attention and motivation, could help restore (and perhaps even enhance) cognitive capabilities.

**Figure 1:**
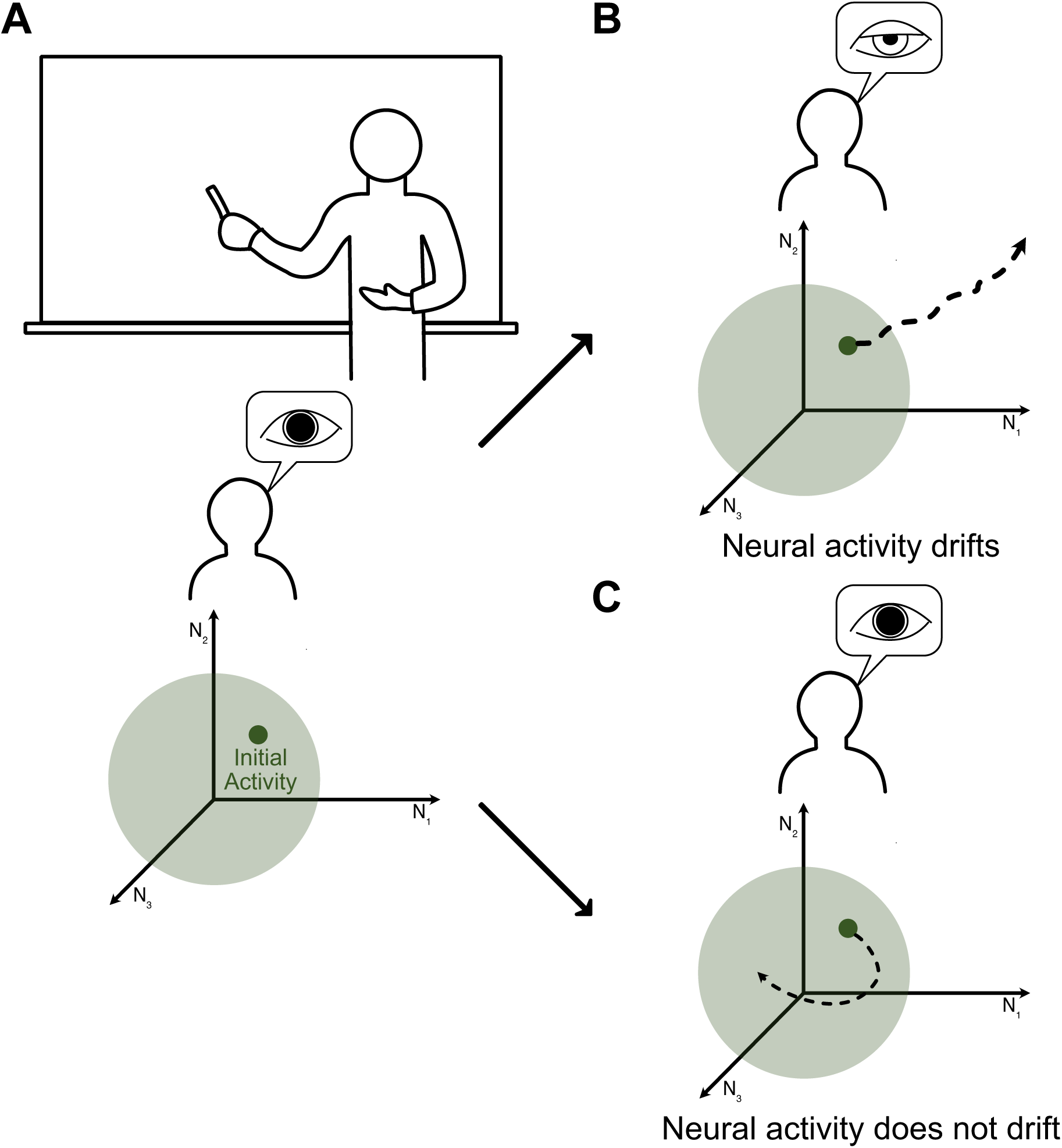
Fluctuations in internal states occur through changes in neural population activ-ity and shape behavior. Changes in internal states, such as arousal, influence sensory processing and behavior. **A.** A student may have high arousal levels at the start of a lecture, as indicated by the large pupil diameter. This may correspond to initial population activity (green dot) located within a region of neural space associated with elevated arousal (green sphere). Each axis of the neural space represents the activity of one neuron (*N_i_* refers to the activity of the *i*-th neuron). **B.** Over time, the student’s arousal may decline, as indicated by a smaller pupil diameter. These changes in internal states manifest as changes in neural population activity, which can drift away from the initial population activity over time (dashed line), potentially exiting the zone associated with elevated arousal. **C.** If this activity drift could be reined in and neural activity could be kept within the region associated with elevated arousal, the student might remain focused on the lecture.

To assess how well animals could volitionally counteract fluctuations in their neural activity over time, we used a brain-computer interface (BCI), which provides moment-to-moment neurofeedback or sensory feedback about neural activity (Orsborn *et al.* [11], Wander *et al.* [12], Moxon *et al.* [13], Golub *et al.* [14], Sitaram *et al.* [15], Watanabe *et al.* [16], Andersen *et al.* [17]). In a BCI, the causal relationship (or “mapping”) between the recorded neural activity and behavior is known exactly and defined by the experimenter. This allows the experimenters to define how animals need to volitionally modify their neural activity for task success. The control signals extracted by a BCI mapping can depend on the brain area involved. When BCIs use activity from motor cortex, mappings often rely on neural activity dominated by movement-related signals. In contrast, activity outside motor cortex may reflect a more heterogeneous mixture of variables that can include movement-related and internal state signals (Rigotti *et al.* [18], Musall *et al.* [19], Steinmetz *et al.* [20]). Previous work has shown that neurofeedback can allow animals to volitionally increase or decrease neural activity in multiple brain areas outside the motor cortex (e.g., Kobayashi *et al.* [21], Shibata *et al.* [22], Clancy *et al.* [23], Hira *et al.* [24], Okazaki *et al.* [25], Sherwood *et al.* [26], MacInnes *et al.* [27], Prsa *et al.* [28], Faller *et al.* [29], Leinders *et al.* [30], Jeon *et al.* [31]). Yet whether animals can use neurofeedback to maintain neural activity outside the motor cortex near an activity target over time, and whether doing so involves regulation of an internal state, remains unknown.

In this work, we trained rhesus monkeys to modulate neural activity in the dorsolateral pre-frontal cortex (PFC) to control the diameter of an annulus on a computer screen via a BCI. PFC has been linked to a range of internal states, such as arousal (Milton *et al.* [32], Cowley *et al.* [5]), attention (Gregoriou *et al.* [33], Snyder *et al.* [34]), motivation (Roesch *et al.* [35]), and decision-related variables (Kiani *et al.* [3]). Animals were required to shrink the annulus, whose diameter reflected how far the animals’ current neural activity was from an activity target. We found that animals kept their neural activity closer to the activity target when visual feedback was driven by concurrently recorded neural activity than during sham trials, in which feedback was decoupled from their ongoing neural activity. Moment-to-moment changes in PFC activity during BCI use were also correlated with pupil diameter changes, suggesting that animals modulated arousal-related internal states during neurofeedback. More broadly, these findings may inform clinical BCIs for restoring impaired internal state regulation and neurofeedback approaches for reducing internal state variability before or during behavioral tasks, allowing researchers to better isolate neural correlates of behavior.

## Results

### A PFC brain-computer interface to self-regulate neural activity

We trained two male adult rhesus macaques to perform a BCI task in which the spiking activity of neurons recorded in PFC drove changes in the diameter of an annulus on a computer screen (Fig. 2A). In this task, the animals’ objective was to shrink the annulus diameter below a pre-specified threshold to get a liquid reward (see Methods). To do so, they had to keep their neural activity as close as possible to an activity target.

**Figure 2:**
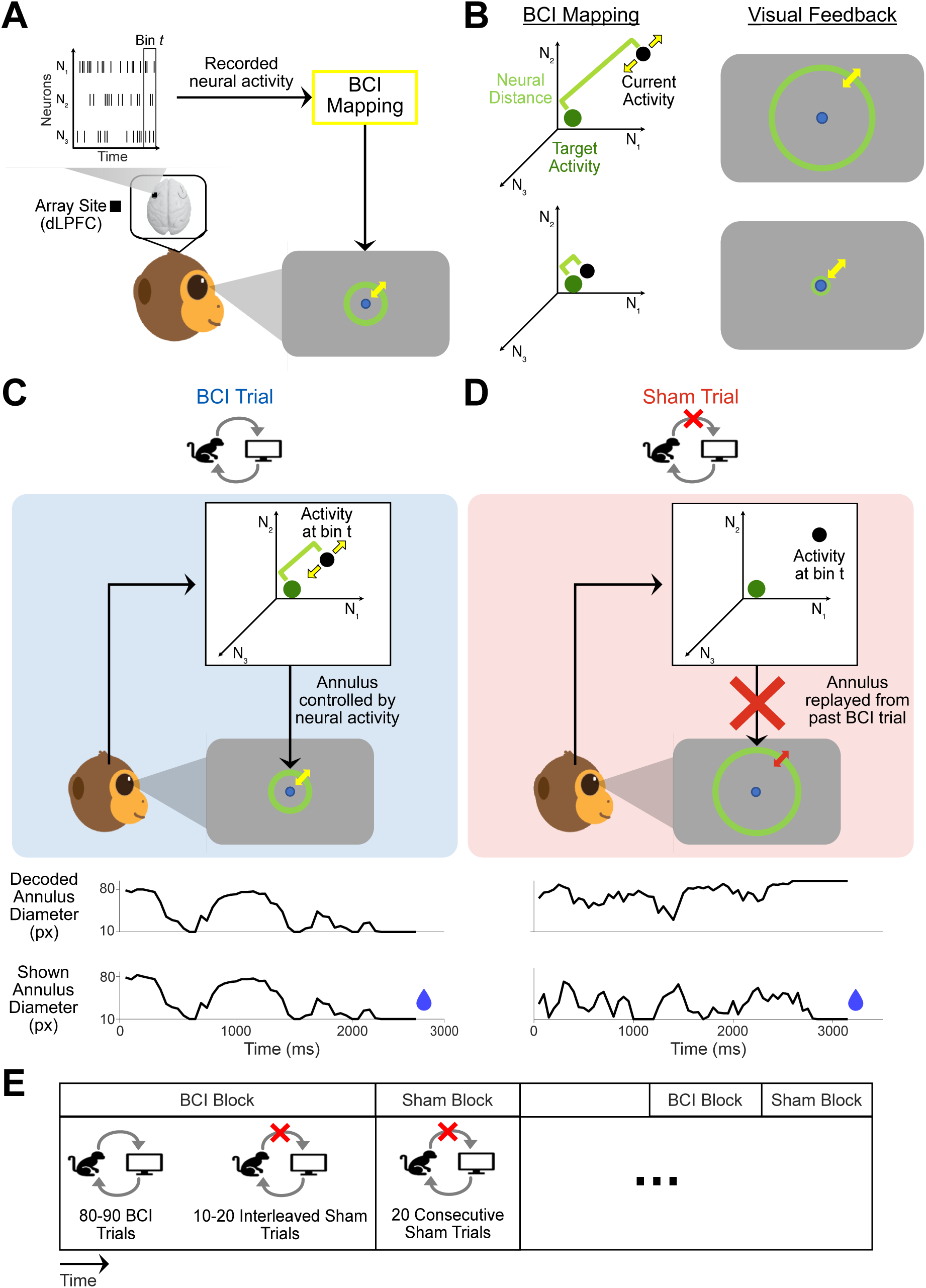
Using a BCI to maintain neural activity near an activity target. **A.** Schematic of BCI in the prefrontal cortex. Neural activity was recorded using a multi-electrode array implanted in the dorsal lateral prefrontal cortex (PFC, black square) and mapped to moment-to-moment visual neurofeedback. **B.** The BCI mapping converted the neural distance (green bracket) from the animals’ current population activity (black dot) to an activity target (green dot) into the diameter of an annulus shown on the screen (yellow arrows denote the expansion and contraction of the annulus). Large neural distances corresponded to large annulus diameters (top row), while small neural distances corresponded to small annulus diameters (bottom row). To succeed, animals had to maintain their neural distances below a threshold set for each day based on the activity observed during calibration. This threshold was chosen such that approximately 50% of the calibration trials would stay below it and be rewarded (see Methods). For illustrative purposes, this schematic shows the neural distance measured in the population activity space, although we computed the neural distance in a latent space in the actual experiments (see Methods). **C.** In BCI trials, the shown annulus diameter was determined by neural activity recorded on the same trial (upper panel). Thus, the decoded and shown annulus diameters (black curves, lower panels) were identical by definition. Liquid reward (blue droplet) was provided when the success criterion was met. **D.** In sham trials, animals viewed replayed feedback during a replay feedback period, in which the annulus diameter was determined by neural activity recorded on a different trial rather than by the animal’s ongoing neural activity (upper panel). Thus, the decoded annulus diameter from the animal’s ongoing neural activity differed from the annulus diameter shown on the screen (see Methods). Liquid reward was provided as long as fixation was maintained throughout the replay feedback period. **E.** Session structure. Animals completed alternating BCI and sham blocks. In most sessions, BCI blocks comprised 90 BCI trials and 10 randomly interleaved sham trials. In a few Monkey S sessions, BCI blocks had 80 BCI trials and 20 interleaved sham trials. Sham blocks comprised 20 consecutive sham trials.

At the start of each experiment, animals completed calibration trials in which they fixated on a gradually shrinking annulus and received rewards for maintaining fixation (see Methods). Neural activity recorded during these trials defined the BCI mapping for the subsequent BCI task. For the BCI mapping, we used factor analysis (FA) to identify latent variables that captured the greatest shared variance among the recorded neurons (Cunningham *et al.* [36]). We defined our activity target as the mean latent activity over time observed during these calibration trials. Thus, the activity target was a 4- or 5-element vector corresponding to the 4 or 5 dimensions of the latent space (see Methods). We did so to assess how proficiently animals could use neurofeedback to prevent drift in PFC population activity, which has previously been linked to fluctuations in the animals’ arousal-related internal states over time (Cowley *et al.* [5]).

After completing the calibration trials, animals performed BCI trials in which annulus diameter provided accurate moment-to-moment neurofeedback on how far the animals’ neural population activity (a spike count vector from the neural population in a 50 ms time bin projected into the 4-or 5-dimensional latent space) was from the activity target (Fig. 2B; see Methods and Fig. S1). A large annulus diameter indicated that the neural activity was far away from the activity target, whereas a small annulus diameter indicated that the neural activity was close to the activity target. To quantify this, we used the *neural distance* (green bracket in Fig. 2B; see Methods), defined as the Euclidean distance between the animals’ current population activity and the activity target in the latent space identified by FA. In BCI trials, the neural distance at the current time bin (bin size of 50ms) directly determined the annulus diameter shown in the subsequent time bin (Fig. 2C). To succeed, animals had to maintain their neural distances below a predetermined threshold for 8 consecutive time bins, at which point the trial terminated with a liquid reward (see Methods).

Notably, animals received no additional instructions and had to use only the annulus feedback to guide their neural activity toward the activity target. This differs from BCIs that use activity from motor cortex, where control signals often rely on directional movement intent (Shenoy *et al.* [37]). Unlike motor cortex, PFC encodes a diverse array of signals related to cognitive control, internal states, sensory information, and motor plans (Miller *et al.* [38]), with no known dominant signal for animals to leverage for volitional control. Thus, our BCI required animals to attain an activity target observed at the start of a session by relying solely on visual feedback of their neural activity.

To evaluate the efficacy of neurofeedback, we also included sham trials, which replayed feedback from the correct BCI trials of previous sessions regardless of the current neural activity, thereby dissociating the visual feedback from the animals’ current neural activity (Fig. 2D, see Methods). These sham trials were important for demonstrating that neurofeedback is more useful to the animals if it is informative about their current neural activity (Caria *et al.* [39], Thibault *et al.* [40], Schneider *et al.* [41], Stefano Filho *et al.* [42]). To encourage animals to stay engaged during sham trials, which always lasted the maximum possible BCI feedback duration of 3.4 seconds, animals received a liquid reward for maintaining fixation throughout the replay feedback period. We used two types of sham trials with complementary roles. Consecutive sham trials were organized in homogeneous blocks and provided a reference success rate for task performance under replayed feedback. During these blocks, there was low incentive for active neural regulation, since reduced neural distances could not shorten the legnth of each trial (Fig. 2E, “Sham Block”). By contrast, interleaved sham trials were randomly interspersed among BCI trials. Because they occurred randomly, the animals’ motivation to achieve low neural distances was presumably maintained (Fig. 2E, “BCI Block”). Together, these sham trials defined a chance level that allowed us to assess whether the neurofeedback was beneficial in reducing neural distances.

### Neural activity during neurofeedback stayed closer to the activity target than with sham feedback

We first asked whether animals could use the BCI to volitionally maintain their neural activity near the activity target. To assess whether neurofeedback enabled above-chance performance, we compared BCI trials to consecutive sham trials. We evaluated the performance of sham trials by processing them in the same way as BCI trials offline. Specifically, we passed the recorded neural activity during sham trials through the BCI mapping and assessed whether the neural activity would have met the success criterion (see Methods and Fig. S2).

We observed that animals maintained their neural activity near the activity target more effectively with neurofeedback, as BCI trials exhibited a higher success rate than consecutive sham trials (Fig. 3A, black line is higher during blue blocks than in red blocks). Task success was defined as a binary outcome on each trial indicating whether the animals met the task criterion, and success rate was computed as the fraction of successful trials within each block of trials. To capture in more detail how well the animals kept their neural activity near the activity target, we also quantified neural distance as a continuous-valued measure. We found that BCI trials had lower per-block average neural distances (averaged across bins within each trial, then across trials within each block) than consecutive sham trials within the same session (Fig. 3B, blue lines are lower than the red lines). Across sessions, we observed a similar trend as BCI trials had higher success rates (Fig. 3C, points below diagonal) and lower neural distances (Fig. 3D, points above diagonal) than consecutive sham trials. These differences indicate that animals could successfully use the BCI to maintain their neural activity near the activity target.

**Figure 3:**
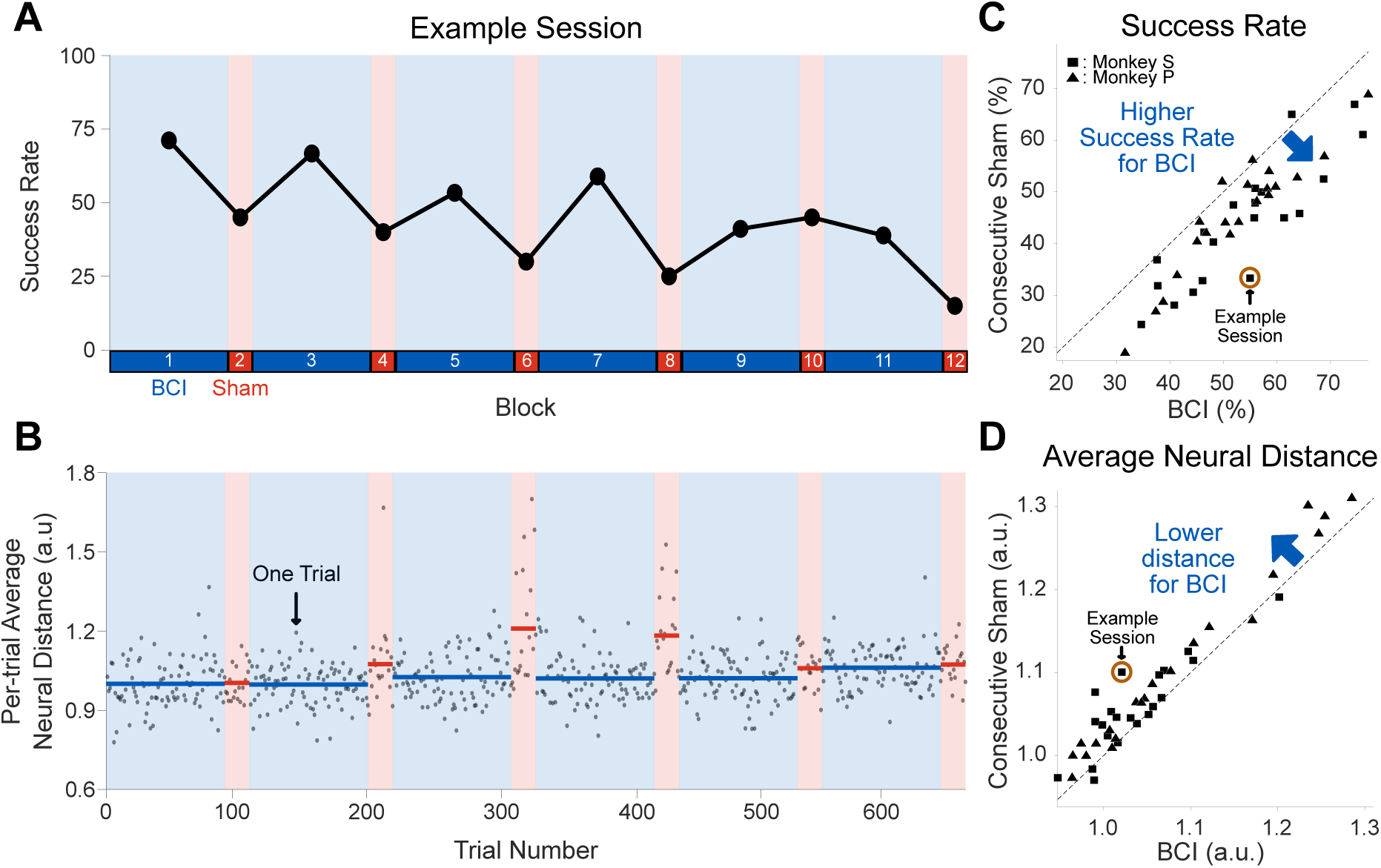
Animals successfully used the BCI to counteract fluctuations in their neural activity. **A.** Success rate in each block of trials for an example session from Monkey S (computed offline for sham blocks; see Methods). The background shading denotes the two block types: BCI (blue) and sham (red). Interleaved sham trials were not included in the success rate calculation of BCI Blocks. **B.** The per-trial average neural distance (averaged across time bins within each trial) for each trial (gray dots) from the same example session as in **A**. Blue and red lines indicate the per-block average neural distances (averaged across trials within each block). **C.** BCI trials had a higher success rate than consecutive sham trials across sessions (one-sided Wilcoxon paired signed rank test; *p <* 10*^−^*^8^ pooled across monkeys, *N* = 42 sessions; *p <* 10*^−^*^4^ for each monkey, N = 21 sessions for each monkey). Success rates were computed for consecutive sham trials after they were processed like BCI trials (see Methods and Fig. S2). Each symbol represents the success rate computed as the number of correct trials divided by the total number of trials for each type in a session. The brown circle indicates the example session shown in panels **A** and **B**. **D.** Per-condition average neural distances (averaged across all trials of a given type within each session) were lower for BCI trials than consecutive sham trials (one-sided Wilcoxon paired signed rank test; *p <* 10*^−^*^6^ pooled across monkeys, *N* = 42 sessions; *p <* 0.005 for each monkey, *N* = 21 sessions for each monkey). Each symbol represents the average neural distance for a session. When computing per-condition average neural distances for consecutive sham trials, we included only time bins retained by the truncation process described in Fig. S2 (see Methods). The brown circle indicates the example session shown in panels **A** and **B**.

### Neural fluctuations away from the activity target were gradually reined in across neurofeedback trials

Slow changes in internal states (Stringer *et al.* [7], Allen *et al.* [8], Cowley *et al.* [5]), such as arousal, can detrimentally impact BCI task performance as they may cause the recorded neural activity to move away from the activity target. We thus wondered if animals improved their performance in BCI blocks by using neurofeedback to counteract slowly varying fluctuations in neural population activity, possibly induced by such internal state changes, that would otherwise cause it to drift away from the activity target across trials within each block.

To investigate whether animals counteracted this drift, we examined how the per-trial average neural distance changed across trials within blocks. We found that sham blocks tended to exhibit gradual increases in neural distance over the course of 60-120 seconds, indicating drift away from the activity target (Fig. 4A, positive slope of red dashed line). However, when analyzing duration-matched segments of BCI blocks and considering only the BCI trials within each segment (see Methods), we found that animals exhibited a smaller increase in per-trial average distances across trials (Fig. 4A, flat slope of blue dashed line), suggesting that animals used neurofeedback to mitigate drift away from the activity target. Across sessions, BCI blocks consistently had lower regression slopes than the sham blocks that immediately followed them (Fig. 4B, points above diagonal), indicating that animals used neurofeedback to either mitigate distance-related drift (near-zero slopes) or even counteract it (negative slopes). However, BCI blocks only partially counteracted sham-related drift, yielding a staircase-like pattern across blocks within a session. Average neural distance tended to increase during sham blocks, and BCI blocks either flattened or partially reversed this increase (Fig. S3, consistent with the gradual performance decline across blocks in Fig. 3A). These results indicate that animals used neurofeedback to counteract drift away from the activity target during BCI trials, mitigating but not fully reversing the drift that accumulated over tens of seconds.

**Figure 4:**
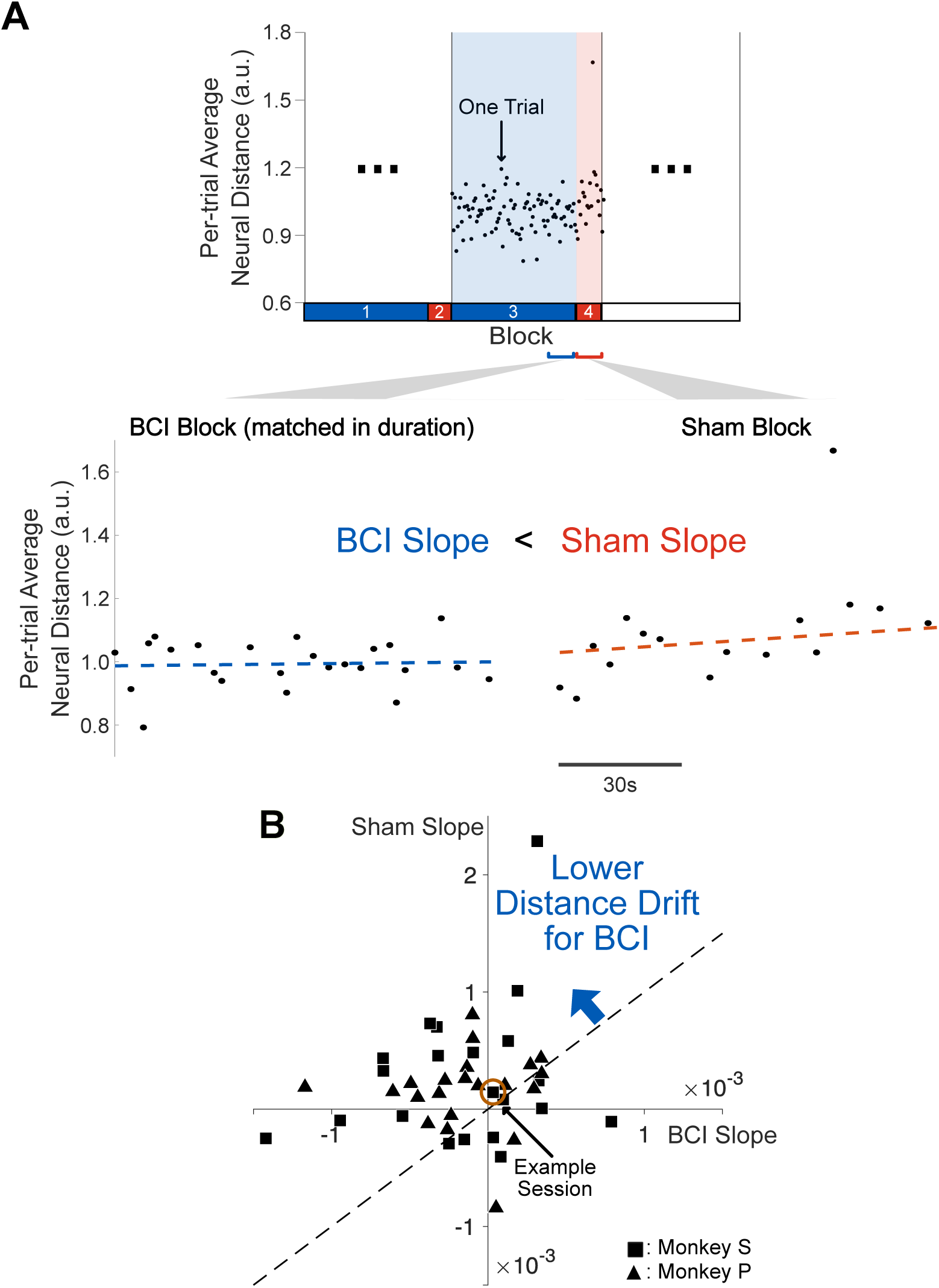
Neural distance increased over the course of sham blocks but not BCI blocks. **A.** In an example session, a BCI block showed a smaller increase in per-trial average neural distance over trials (dashed blue line, linear regression slope = 1.2 × 10*^−^*^4^) compared to the subsequent sham block (dashed red line, slope = 8.0 × 10*^−^*^4^). BCI blocks were truncated to match the subsequent sham block in clock time duration (in seconds). Interleaved sham trials were not included in the slope calculation for BCI blocks. A positive slope indicates that neural activity drifted away from the activity target over time, whereas a negative slope indicates that the animals brought their neural activity closer to the activity target over time. **B.** Slopes for BCI blocks were significantly lower than the slopes for sham blocks (one-sided Wilcoxon paired signed rank test; *p <* 10*^−^*^6^ pooled across monkeys, *N* = 42 sessions; *p <* 0.05 for each monkey, *N* = 21 sessions for each monkey). Each symbol represents the average regression slope computed across blocks within a session, after the regression slope was found separately for each block. For sham blocks, we computed regression slopes using the per-trial distances computed from the time bins retained by the truncation process described in Fig. S2 (see Methods). The brown circle indicates the example session shown in **A**. The regression slope for sham blocks within a session was positive (median of 1.94 × 10*^−^*^4^ for sessions pooled from both monkeys; median of 1.45 × 10*^−^*^4^ for monkey S and 2.01 × 10*^−^*^4^ for monkey P; one-sided Wilcoxon signed rank test; *p <* 0.005 pooled across monkeys, *N* = 42 sessions; *p <* 0.05 for each monkey, *N* = 21 sessions for each monkey). In contrast, the regression slope for BCI blocks within a session was negative (median of −1.17 × 10*^−^*^4^ for sessions pooled from both monkeys; median of −9.42 × 10*^−^*^5^ for monkey S and −1.34 × 10*^−^*^4^ for monkey P; one-sided Wilcoxon signed rank test; *p* = 0.0208 pooled across monkeys, *N* = 42 sessions; *p* = 0.1119 for monkey S and *p* = 0.0549 for monkey P, *N* = 21 sessions for each monkey). This indicates that sham blocks exhibited a gradual distance drift while BCI blocks did not.

### Animals leveraged moment-to-moment visual feedback to counteract neural fluctuations

Our previous analyses demonstrated that animals could successfully use the BCI to maintain their neural activity near the activity target (Fig. 3) and counteract gradual fluctuations away from it (Fig. 4). However, internal states can also fluctuate on faster timescales of a few seconds (Cohen *et al.* [2], McGinley *et al.* [4], Hennig *et al.* [6]), raising the question of whether animals could also use moment-to-moment visual details of neurofeedback to counteract within-trial fluctuations in neural activity. The comparison between BCI and consecutive sham trials does not adequately address this question, as the observed differences (Fig. 3 and Fig. 4) might have reflected animals adopting different strategies across block types. For instance, rather than having actively used the moment-to-moment visual feedback, animals may have instead recognized that BCI trials ended early upon reaching the activity target, whereas consecutive sham trials always ran for the full feedback period. This difference may have led them to adjust their attempts to regulate their neural activity accordingly.

To directly test whether animals used moment-to-moment neurofeedback rather than block-level strategies to counteract these fluctuations within a trial, we examined sham trials that were randomly distributed within BCI blocks (Fig. 5A). Unlike consecutive sham trials, in which all trials within a block were sham trials, interleaved sham trials were rare within BCI blocks and occurred amongst the BCI trials, with animals therefore likely anticipating valid neurofeedback. If animals relied on moment-to-moment visual details of neurofeedback to keep their neural activity close to the activity target, BCI trials would show higher task performance compared to interleaved sham trials.

**Figure 5:**
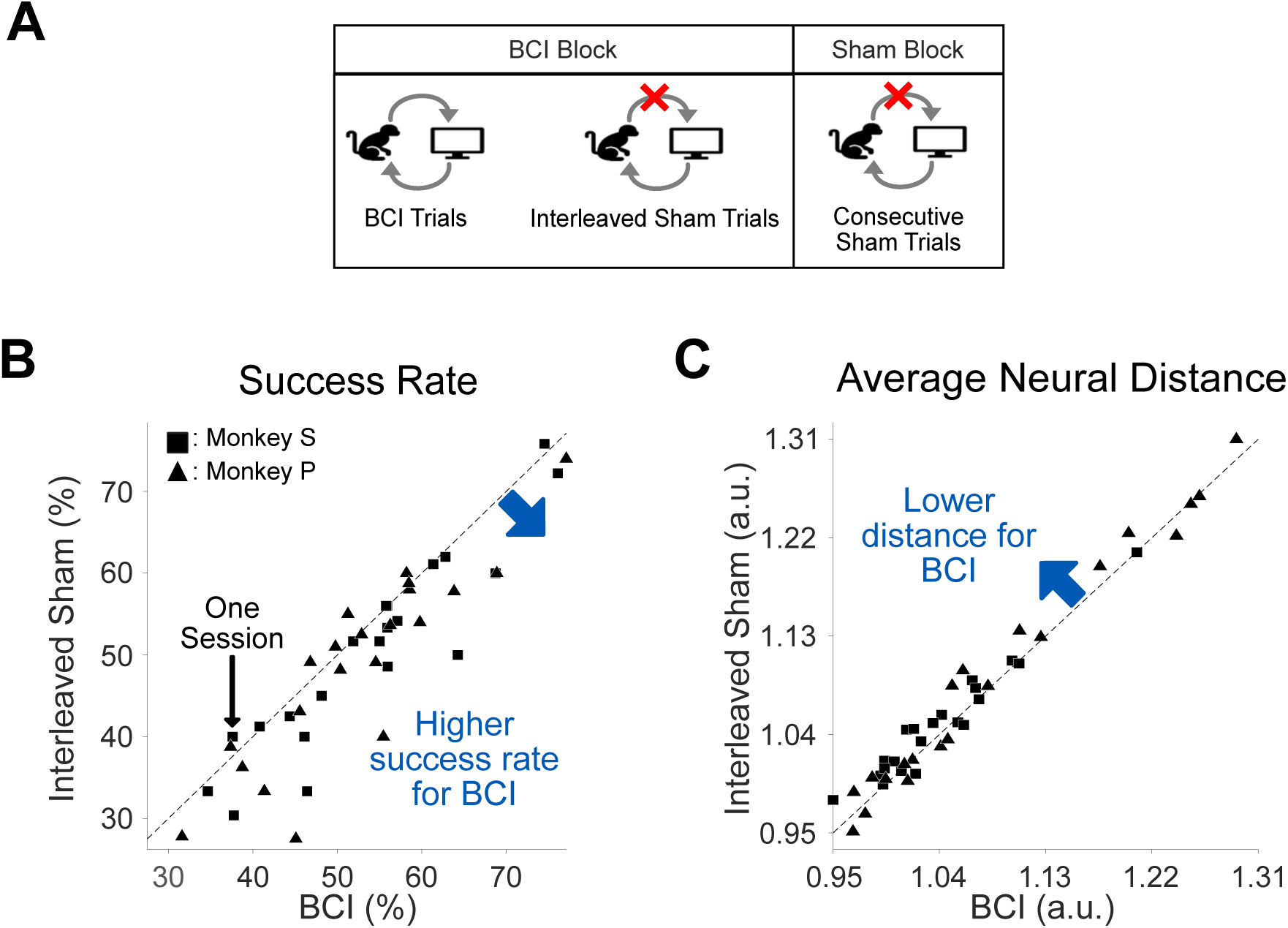
Animals used moment-to-moment neurofeedback to perform better in BCI trials than in interleaved sham trials. **A.** We used two types of sham trials: interleaved sham and consecutive sham trials. Interleaved sham trials, unlike consecutive sham trials, were presented within the same blocks as BCI trials. **B.** Success rate was higher for BCI trials than for the interleaved sham trials (one-sided Wilcoxon paired signed rank test; *p <* 10*^−^*^4^ pooled across monkeys, *N* = 42 sessions; *p <* 0.05 for each monkey, *N* = 21 sessions for each monkey). Success rates were computed for interleaved sham trials after they were processed like BCI trials (see Methods and Fig. S2). Each symbol represents the success rate computed as the number of correct trials divided by the total number of trials for each type in a session. **C.** Per-condition average neural distances were lower for BCI trials than interleaved sham trials (one-sided Wilcoxon paired signed rank test; *p* = 0.0023 pooled across monkeys, *N* = 42 sessions; *p* = 0.0051 for monkey S and *p* = 0.0675 for monkey P; *N* = 21 sessions for each monkey). When computing per-condition average neural distances for interleaved sham trials, we only included time bins retained by the truncation process described in Fig. S2 (see Methods). Each symbol represents the average neural distance for a session.

We evaluated interleaved sham trial performance by processing these trials offline in the same way as BCI trials (see Methods and Fig. S2). We found that animals exhibited higher success rate (Fig. 5B, points below diagonal) and smaller average neural distances (Fig. 5C, points above diagonal) in BCI trials than interleaved sham trials. This indicates animals used neurofeedback on a moment-to-moment basis to keep their neural activity close to the activity target. Furthermore, we discovered that the context in which sham trials appeared mattered, as interleaved sham trials had higher success rates and lower average neural distances than consecutive sham trials (Fig. S4). This result suggests that animals kept trying to reduce their neural distances in interleaved sham trials even when shown feedback did not reflect their current neural activity, possibly by using an internal model of the BCI (Golub *et al.* [43]). Taken together, these findings indicate that animals performed better in BCI than in sham trials by using the moment-to-moment visual details of neurofeedback to regulate their neural activity on a timescale of hundreds of milliseconds.

### BCI use was associated with arousal-related changes in neural activity

Given that animals improved BCI performance by counteracting neural activity fluctuations over seconds, we sought to identify the mechanisms underlying this regulation. Because internal states such as arousal fluctuate over similar timescales and can covary with PFC activity (Milton *et al.* [32], Cowley *et al.* [5]), we asked whether arousal-related changes were associated with BCI use. We first examined whether recorded PFC activity tracked pupil diameter, a widely used measure of arousal (Joshi *et al.* [44]), and found a slowly varying pupil-linked arousal signal in PFC activity (Fig. S5). One possible source of this signal is the locus coeruleus (LC; Fig. 6A, green circle). The LC mediates arousal through widespread norepinephrine (NE) release to downstream areas, including PFC, and its activity covaries with pupil diameter (Sara [45], Joshi *et al.* [44]). Thus, if LC-mediated arousal fluctuations contributed to PFC activity during BCI use, annulus diameter, which was driven by PFC activity, should have covaried with pupil diameter across all time bins within a session, capturing both within-trial and across-trial fluctuations (Fig. 6A).

**Figure 6:**
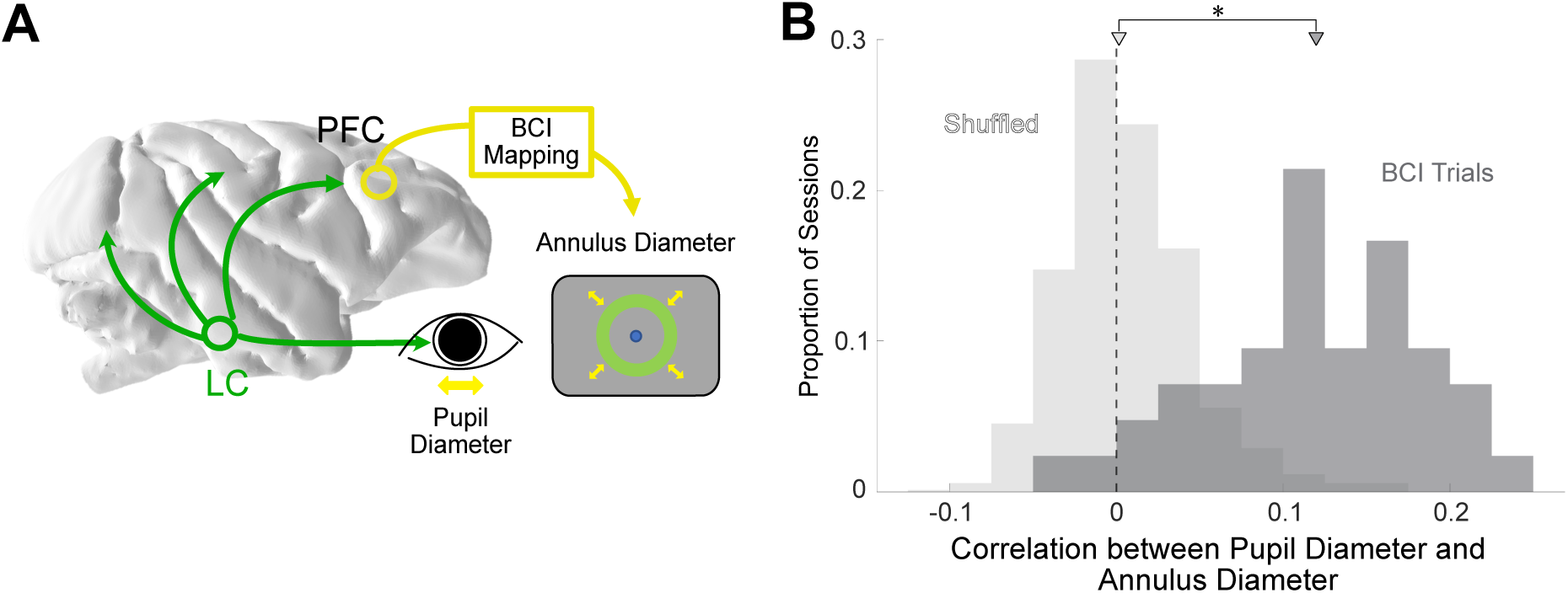
Pupil-linked fluctuations in arousal tracked BCI performance. **A.** Conceptual illustration of how animals may modulate their neural activity over time by regulating an arousal-related internal state. The Locus Coeruleus (LC, green circle) projects to the PFC and is involved in regulating arousal. Green arrows emanating from the LC indicate its projections throughout the cortex. The LC can also modulate arousal-related pupil diameter changes (green arrow leading to the eye). Animals may have regulated an arousal-related internal state to use the BCI (yellow arrow from PFC to BCI mapping). **B.** Pupil-linked arousal fluctuations tracked the annulus diameter during BCI trials. Pupil diameter was correlated with the shown annulus diameter on a bin-by-bin basis (bin size of 50ms) during BCI trials. For each session, we computed the Pearson correlation coefficient between the pupil diameter and the shown annulus diameter for all bins in BCI trials (median Pearson correlation coefficient of *r* = 0.1212 for sessions pooled from both monkeys). For comparison, we computed a chance distribution by shuffling the data across sessions–we paired pupil data from one session with annulus diameters from different sessions–before computing the Pearson correlation between pupil and annulus diameter (median *r* = 0.0020 for sessions pooled from both monkeys; see Methods). The correlation was significantly higher (one-sided Wilcoxon rank sum test; *p <* 10*^−^*^19^ pooled across monkeys, *N* = 42 sessions; *p <* 10*^−^*^9^ for each monkey, *N* = 21 sessions for each monkey) for the observed data (dark gray histogram) than for the shuffled data (light gray histogram). Triangles indicate the median of each distribution.

We found that during BCI trials, the annulus diameter was indeed positively correlated with the pupil diameter of the animals across all time bins within a session (Fig. 6B and see Methods). We focused only on BCI trials because in those trials animals received moment-to-moment feedback about their PFC activity, which may have reflected changes in their pupil-linked internal states. The positive correlations show that higher BCI performance (i.e., smaller neural distances) was associated with smaller pupil size, consistent with maintaining a more stable and lower level of arousal. However, a potential confound is that smaller annulus diameters would concentrate more luminance on the fovea (though the smaller annulus area would mean less total luminance from the screen), which could induce pupil diameter decreases and lead to the observed positive correlation. We performed a control analysis to show that these pupil-annulus correlations were unlikely to be driven solely by luminance-related pupillary responses (Fig. S6). Overall, these results demonstrate that one mechanism by which animals achieved BCI success was through the self-regulation of an internal state linked to arousal.

## Discussion

In this work, animals used a BCI to maintain their PFC neural activity close to an activity target. To improve their performance, animals relied on neurofeedback to volitionally counteract activity fluctuations that occurred over timescales of seconds to hundreds of milliseconds. Furthermore, we found that self-regulation of neural activity coincided with changes linked to arousal. Taken together, our findings suggest that animals can use neurofeedback to volitionally regulate their neural activity by modulating their arousal levels.

Our work exemplifies how neural activity outside of motor cortex can be brought under volitional control using a BCI. Prior work has demonstrated volitional regulation of neural activity in several regions outside motor cortex using BCIs, including visual cortex (Shibata *et al.* [22], Neely *et al.* [46], Schneider *et al.* [41], Clancy *et al.* [47], Jeon *et al.* [31], Chinchani *et al.* [48]), somatosensory cortex (Clancy *et al.* [23]), parietal cortex (Bagherzadeh *et al.* [49]), prefrontal cortex (Kobayashi *et al.* [21], Schafer *et al.* [50], Sherwood *et al.* [26], Leinders *et al.* [30]), ventral tegmental area (MacInnes *et al.* [27]), and insula (Caria *et al.* [39]). Although some of these studies include demonstrations of BCI-based volitional control of PFC neural activity, PFC remains a challenging target for BCI use because it encodes diverse signals related to cognitive control, internal states, sensory information, and motor plans (Miller *et al.* [38]). Despite this complexity, our findings expand understanding of volitional control outside motor cortex by showing that animals can volitionally regulate PFC activity through modulation of an arousal-related internal state signal.

Changes in internal states can affect many neurons, and population-level signatures of internal states have been inferred by relating neural activity to behavioral or physiological measures (Allen *et al.* [8], Stringer *et al.* [7], Cowley *et al.* [5]). Dimensionality reduction techniques, such as factor analysis, can identify these population activity patterns reflecting shared neural variability, which can then be linked to internal states using measurements such as pupil size. Prior intracortical BCIs using activity outside motor cortex have used mappings based on single-electrode activity (Kobayashi *et al.* [21], Schafer *et al.* [50]) or activity from two predefined groups of neurons (Neely *et al.* [46]). Although not explicitly designed to regulate internal states, such mappings may be limited in their ability to infer distributed population-level signals. Our findings support the feasibility of population-level approaches for future BCIs aimed at internal state regulation.

Our work demonstrates that control of a PFC BCI may engage neuromodulatory circuits involved in mediating arousal. One plausible mechanism is that the regulation of PFC activity during BCI use modulates the LC, which mediates NE release and also influences pupil diameter (Fig. 6). However, pupil diameter is not uniquely affected by the LC-NE system and can also reflect other internal states, including attention, emotion, or belief uncertainty (Joshi *et al.* [51]). These states may be mediated by additional neuromodulatory systems, such as those involving acetylcholine (ACh) and serotonin (5-HT). Given PFC’s connections with the basal forebrain and dorsal raphe nuclei, which are the major cortical sources of ACh and 5-HT, respectively (Lin *et al.* [52], Puig *et al.* [53]), these systems may also contribute to pupil-linked changes during BCI control. Thus, while our findings suggest that PFC BCI control may involve the regulation of an arousal-related internal state, further studies combining BCI use with direct recordings or perturbations of neuro-modulatory nuclei are needed to identify which mechanisms are recruited during BCI control.

Our BCI paradigm helps experimenters reduce internal state variability, a source of trial-to-trial variability that can obscure links between neural activity and behavior. Even under identical task conditions, animals’ internal states can change over time, affecting behavior outside of the experimenter’s control. For example, in perceptual decision-making tasks, animals may enter ‘lapse’ states and make errors despite strong sensory evidence favoring one choice over the other (Ash-wood *et al.* [54]). In stimulus change discrimination tasks, animals may also vary in impulsivity, reporting which stimulus changed before fully integrating the available sensory evidence (Cowley *et al.* [5]). Our neurofeedback framework provides a way to rein in such internal state-related neural variability on a trial-by-trial (i.e., seconds) and moment-by-moment (i.e., hundreds of milliseconds) basis. In each trial, animals could first use neurofeedback to attain a desired internal state before performing the behavioral task, providing a cleaner readout of the neural correlates of behavior (Motiwala *et al.* [55]).

BCIs, such as the one presented here, have the potential to help users regulate internal states, with broad implications for clinical applications (Sterman *et al.* [56], Loriette *et al.* [57]). Dysregulation of internal states contributes to many neuropsychiatric and emotional disorders, including attention-deficit hyperactivity disorder and post-traumatic stress disorder (Sonuga-Barke *et al.* [58], Sani *et al.* [59], Kredlow *et al.* [60]). Such dysregulation may also affect clinical BCIs designed to restore movement, in which fluctuating internal state signals in the primary motor cortex can interfere with motor intention signals and degrade BCI performance (Gallego *et al.* [61]). Because many internal states, including attention (Cohen *et al.* [2], Snyder *et al.* [34], Chinchani *et al.* [48]), mood (Sani *et al.* [59]), and motivation (Smoulder *et al.* [62]), can be decoded from neural population activity, our neurofeedback paradigm offers a way to guide population activity patterns toward those associated with desired internal states. With repeated neurofeedback use, training could progressively reinforce neural activity linked to these desired internal states (Athalye *et al.* [63]) and alter the neural activity that is produced even when neurofeedback is no longer used (Losey *et al.* [64])

## Methods

### Experimental procedures

All experimental procedures were approved by the Institutional Animal Care and Use Committees of the University of Pittsburgh and Carnegie Mellon University. Two adult male rhesus macaque monkeys (Monkey **S**: 8 years; Monkey **P**: 10 years) were used for this study. Surgeries on animals were performed in sterile conditions under general anesthesia using isoflurane.

### Neural and behavioral recordings

We recorded from the prefrontal cortex on the gyrus anterior to the arcuate sulcus (area 8Ar of the dorsolateral prefrontal cortex) with a 96-electrode Utah array (10x10, 1.0 mm, Blackrock Microsystems). We recorded from the right hemisphere in Monkey **S** and the left hemisphere for Monkey **P** while they sat head-fixed in a primate chair. Data collection was performed using custom software in Matlab (Mathworks) that utilized Psychophysics toolbox extensions (Kleiner *et al.* [65]). Visual stimuli were displayed on a 21” cathode ray tube (CRT) monitor at a resolution of 1024×768 pixels and a refresh rate of 100 Hz. The animals’ eyes were 36 cm from the screen. Signals were band-pass filtered (0.3 - 7500 Hz) and then digitized at 30kHz before being stored offline for analysis. For each electrode, spiking waveforms were defined as a 52-sample (1.73 ms) window of the filtered voltage signal triggered by the signal crossing a predefined threshold. The threshold was defined as four times the root-mean-square voltage of the raw signal on each electrode recorded at the beginning of the session. All behavioral and neural data were recorded using a Grapevine recording system (Ripple Neuro, Millcreek, Utah). Pupil diameter was monitored monocularly at a rate of 1kHz using infrared eye-tracking software (EyeLink 1000; SR Research, Ottawa, Ontario). To isolate waveforms related to neural spiking, we applied a neural network classifier we previously developed that labeled recorded spike waveforms as “neural” or “not neural” (Issar *et al.* [66]). During the BCI experiment, this preprocessing step was applied online to calibration trials and BCI trials. It was also applied offline to sham trials for subsequent offline analyses. Spikes on a given channel were identified as threshold crossings and were counted in non-overlapping 50ms bins. Throughout this work, we refer to each channel as a ‘neural unit’ and the set of spike counts identified in all channels during a 50ms bin as a ‘spike count vector’. We recorded from 35.24 ± 1.95 (mean ± 1 s.d.) neural units across 21 sessions for Monkey S, and 32.86 ± 10.28 neural units across 21 sessions for Monkey P.

### PFC BCI design

We sought to understand whether animals could use a BCI to volitionally counteract fluctuations in their PFC neural activity. To address this question, we designed a BCI task in which the animals had to keep their neural activity close to an activity target. To ultimately develop BCIs for regulating internal states independently of other cognitive tasks, we first asked whether PFC activity could be volitionally controlled without engaging specific cognitive processes, such as those involved in working memory. To do so, we sought to identify an activity target in the absence of any specific cognitive demands. In addition, we wanted the visual feedback during the identification of the activity target to approximate the visual feedback during BCI control. Hence, we structured our experiments so that animals first completed a set of calibration trials, which required them to maintain passive fixation while viewing an annulus whose diameter changed automatically (see details below). We then defined a BCI mapping and an activity target using the neural activity recorded in these trials. During the BCI task, animals were provided with moment-to-moment visual feedback of their neural activity using an annulus. Their goal was to keep their neural activity close to the activity target.

We chose to use an annulus with a variable diameter to provide visual feedback of the neural activity. Since some PFC neurons exhibit selectivity for visual motion direction (Zaksas *et al.* [67], Kim *et al.* [68]), center-out BCI cursor movements, such as those used in BCIs involving the motor cortex, may drive direction-specific visual responses that could potentially interfere with volitional changes in neural activity. The annulus design mitigated this issue by providing neurofeedback through diameter changes that avoided directional biases. The variable diameter provided one-dimensional BCI feedback, a paradigm commonly used in other non-motor BCIs (e.g., Schafer *et al.* [50], Shibata *et al.* [22], Neely *et al.* [46]). Lastly, to ensure that the neurofeedback itself did not evoke visual responses from the PFC neurons involved in BCI usage, we displayed the annulus in a region of the computer screen that was largely outside of regions that induced visual responses in the PFC neurons (determined by a receptive field mapping performed after the array implant). In addition to BCI trials, animals completed sham trials in which they were provided with visual feedback that did not reflect their current neural activity. BCI trials were compared with sham trials to assess the efficacy of neurofeedback in reducing fluctuations in PFC neural activity. We included two types of sham trials: 1) consecutive sham trials, which were organized in homogeneous blocks and permitted a direct comparison between blocks of sustained BCI feedback and its absence and 2) interleaved sham trials, which were randomly interspersed among BCI trials to maintain animals’ motivation to regulate their neural activity, and allowed us to assess whether the animals used the moment-to-moment feedback of their neural activity.

### Experimental Session Structure

To determine the BCI mapping, we first had animals complete a calibration task consisting of 60 calibration trials (see details below). Following the calibration task, the animals were presented with alternating BCI blocks and sham blocks of trials (Fig. 2E). BCI blocks were 100 trials long and contained primarily BCI trials with randomly interleaved sham trials (10% of the 100 trials were sham trials for both monkeys, except we used 20% sham trials in 6 sessions for Monkey S). Sham blocks contained 20 consecutive sham trials. We only analyzed BCI and sham blocks that had all trials in the block completed, as sometimes animals reached satiation at the end of a session before completing all trials in the block. Data were collected across 21 sessions each for Monkey S and Monkey P.

### BCI calibration trials

In a calibration trial, a fixation dot (0.59 degree diameter) was displayed at the center of the screen against an isoluminant gray background. A trial was initiated when the animals fixated on the dot, at which time a green annulus appeared on the screen centered on the fixation dot. The annulus remained fixed in diameter for 400 ms post-fixation, during which the animals were required to maintain fixation. After this fixation period, the annulus started shrinking at a constant speed from a diameter of 80 pixels to a diameter of 10 pixels (a range of 4.71 degrees to 0.59 degrees) during a period of 3.4 seconds. Animals were required to maintain fixation throughout this period and observe the annulus shrink to the diameter that would indicate successful trial completion in the subsequent BCI task. Calibration trials used a shrinking annulus to reinforce the objective of reducing the annulus diameter to obtain a reward in the subsequent BCI task. In addition, the visual stimuli in calibration and BCI trials were designed to be as similar as possible to minimize the extent to which differences in visual input could interfere with BCI performance.

### Defining the BCI mapping

Using the calibration trials, we took spike counts in non-overlapping 50ms bins during the passive fixation period in which the annulus shrank, yielding 68 bins for each trial. We concatenated the binned spike counts across all 60 calibration trials and applied factor analysis (FA) to identify latent variables that capture the greatest variance shared amongst neural units (Cunningham *et al.* [36]). Factor analysis is defined by:

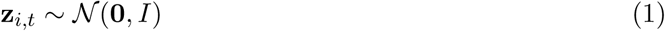

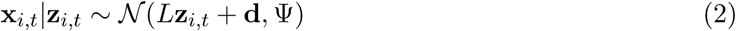

where **z***_i,t_* ∈ ℝ*^m^* are the latent variables at time bin *t* (50ms bins) from calibration trial *i*, **x***_i,t_* ∈ ℝ*^p^* is a vector of binned spike counts across the *p* simultaneously-recorded neural units at time bin *t* of calibration trial *i*, **d** ∈ ℝ*^p^* is a vector of mean spike counts, *L* ∈ ℝ*^p×m^* is the loading matrix relating the latent variables to the neural activity, and Ψ ∈ ℝ*^p×p^* is a diagonal matrix of independent variances for each neuron. The model parameters **d**, *L*, and Ψ were estimated using the expectation-maximization (EM) algorithm.

For consistency, the number of latent variables, *m*, was fixed in all sessions for each animal (Monkey **S**: 5 latent variables and Monkey **P**: 4 latent variables). To validate our choices, we performed a post hoc analysis to identify the appropriate number of latent variables for each session’s calibration trials. For each session, we used FA to identify *d*_shared_, the smallest number of latent variables needed to capture at least 95% of the variability shared amongst the recorded units (Williamson *et al.* [69]). For each monkey, the fixed value of *m* exceeded or was equal to the median of that monkey’s distribution of *d*_shared_ (Monkey **S**: 3 latent variables and Monkey **P**: 4 latent variables), indicating that the chosen number of latent variables was sufficient to capture at least 95% of the variability shared amongst the recorded units across sessions.

We then projected each spike count vector into the *m*-dimensional latent space by computing the posterior mean **ẑ***_i,t_* ∈ ℝ*^m^*:

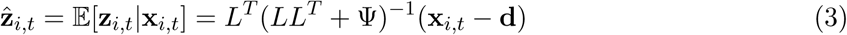

For each calibration trial *i*, we applied exponential smoothing across time bins in the latent space with a smoothing constant *α* ∈ [0, 1] to obtain smoothed latent variables **w***_i,t_* ∈ ℝ*^m^*:

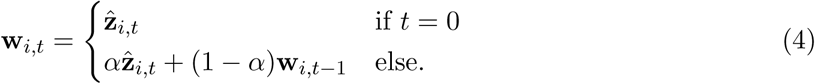

At each bin *t*, the smoothed latent variable **w***_i,t_* was a weighted average of all observed neural activity from calibration trial *i* up to time bin *t* with the weights exponentially decaying over time. The smoothing constant, *α*, thus determines the degree of smoothing (i.e., a smaller *α* corresponds to heavier smoothing/reliance on past neural activity). In all experiments, we used *α* = 0.1535, which corresponded to constant of approximately 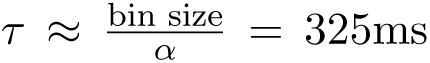, for a bin size of 50ms. This smoothing constant was chosen empirically so that the two criteria were met: 1) there was enough smoothing such that the changes in the neurofeedback over time were not too abrupt, and 2) there was not so much smoothing that it would prevent the neurofeedback from accurately reflecting changes in the animals’ current neural activity. The exponential smoother was reset at the beginning of each calibration trial such that the smoothed latent variable for the first bin of each trial was simply its posterior mean, as indicated in (4).

We defined the activity target ***µ*** ∈ ℝ*^m^* as the mean of these smoothed FA projections:

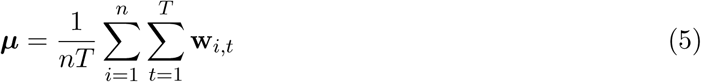

where *n* is the number of calibration trials performed and *T* = 68 refers to the number of 50ms time bins in each calibration trial.

In all sessions, ***µ*** was very close to the origin of the latent space, which corresponds to the mean activity observed for each neural unit **d**. This is due to a property of the FA model in which the mean of the projections for our training data (all bins in our calibration trials) is the zero vector by definition:

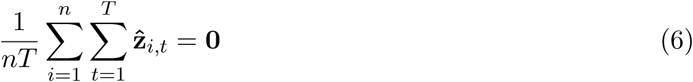

The activity target, ***µ***, slightly deviated from **0** because of the exponential smoothing we applied to the projections before computing the mean.

Next, we used ***µ*** to define the BCI mapping from the recorded neural activity to the neuro-feedback (i.e., annulus diameter) that we provided to the animals (Fig. S1). First, we computed 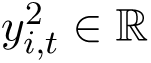, the squared Euclidean distance of the smoothed FA latent variable from the activity target for each time bin *t* of calibration trial *i*:

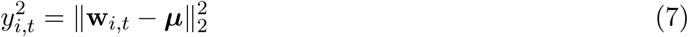

where *t* = 1*, …, T*. Second, to provide neurofeedback-based on the relative rank of the squared neural distances with respect to the calibration dataset, we computed percentiles of *y*^2^ from 0.1 to 99.9, in 0.1 percentile increments from the calibration data. Lastly, we identified the reward threshold, *r*, by sweeping percentile increments from 1 to 99 in steps of 1 percentile. The reward threshold *r* was defined as the percentile increment of the squared distance that would lead to the animals getting approximately 50% of the calibration trials correct when we simulated processing the calibration trials as BCI trials. We selected this 50% success rate to maintain the animals’ motivation to perform the BCI task (Vernon *et al.* [70], Kobayashi *et al.* [21]). For both the simulation and the subsequent BCI task, a trial was considered correct when the animals’ squared neural distances stayed below the reward threshold for 400ms (i.e., 8 consecutive time bins). The reward threshold identified through this process was used for the rest of the trials in that session’s BCI task (Fig. S1B).

During BCI trials, the BCI mapping related the current squared neural distance percentile increments in bin *t* of BCI trial *j* to the next bin’s annulus diameter in pixels, *a_j,t_*_+1_:

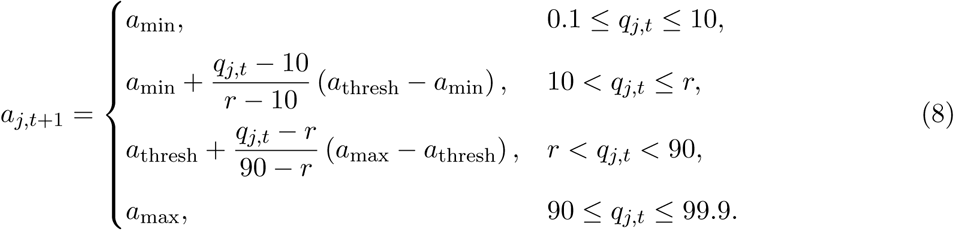

where *q_j,t_*represents the calibration-based percentile closest to the squared neural distance of a given time bin for BCI trial *j*, *a*_min_ = 10px (0.59 degree) was the minimum annulus diameter, *a*_max_ = 80px (4.71 degrees) was the maximum annulus diameter, and *a*_thresh_ = 24px (1.41 degrees) was the annulus diameter when the reward threshold was achieved.

This mapping converts *q_j,t_* into *a_j,t_*_+1_ using a four-segment piecewise-linear mapping: a constant segment at *a*_min_ for percentiles below 10, a linear segment from the 10th percentile to the reward threshold, *r*, a second linear segment from *r* to the 90th percentile, and a constant segment at *a*_max_ for percentiles above 90. (Fig. S1C). This piecewise mapping ensured that *r* always corresponded to the *a*_thresh_, with separate linear slopes for percentile values below and above *r*. As a result, changes in decoded activity were reflected differently depending on whether *q_j,t_* was below or above *r*, providing graded feedback about the animals’ neural activity relative to the success criterion.

### BCI feedback trials

Like calibration trials, BCI trials began with a 400ms fixation period during which the annulus remained still, followed by a 3.4s feedback period. In BCI trials, the animals were given veridical neurofeedback so that the annulus diameter reflected how far their current neural activity was from the activity target. The diameter of the shown annulus at time bin *t* was directly determined by the neural activity observed during the previous *t* − 1 time bins (Fig. S1). First, we projected the spike count vector at bin *t* into the calibration-defined latent space (Eq. 3). Second, we computed the smoothed latent variable (Eq. 4). Third, we computed the squared distance of the smoothed latent variable to ***µ*** (Eq. 7), where ***µ*** was defined using the session’s calibration trials (Eq. 5). Finally, we mapped this squared neural distance to an annulus diameter (Eq. 8). Although the annulus only changed diameters during the feedback period, neural distances were also computed and tracked for bins during the fixation period (i.e., the first 400 ms, or 8 bins, of a trial). This fixation period provided enough time for the exponential smoother (Eq. 4) to stabilize. Thus, when the feedback period started at the 9th bin of a trial, the neural distances calculated for the previous 8 bins were used to determine the initial annulus that was shown to the animals.

During these trials, the animals’ objective was to maintain their squared neural distance below the reward threshold (i.e., keeping the annulus diameter below 24 pixels) for 400ms (Fig. 2C). If the animals maintained their squared neural distances below the reward threshold for 400ms (8 consecutive bins), the trial would end with the animals receiving a liquid reward. If the animals did not achieve this objective during the feedback period, the trial ended with no reward delivery. We selected a 3.4s feedback period to provide sufficient time to try using the BCI feedback, while ensuring the duration was not so long as to lead to disengagment from the task.

### Sham feedback trials

The description that follows applies to both interleaved and consecutive sham trials. In sham trials, the animals were given visual feedback that was not related to how far their current neural activity was from the activity target. Instead, they were replayed feedback from correct BCI trials of previous sessions. During these trials, the animals’ objective was to maintain fixation throughout the replay feedback period. If they maintained fixation for the full duration of the replay feedback period, they received a liquid reward. After the initial 400ms fixation period (identical to what occurred during calibration and BCI trials), animals viewed replay feedback, which was the annulus time course observed during a randomly chosen correct BCI trial of a previous session. The visual feedback during these sham trials was thus unaffected by the animals’ current neural activity. As long as the animals maintained fixation throughout the replay feedback, they were given a liquid reward. To ensure the animals did not break the association between a shrinking annulus and receiving a reward, we specifically selected feedback from correct BCI trials in which the animals met the success criterion at the very end (i.e., exactly at the 68th bin) of the 3.4s feedback period. At the beginning of each session, we determined a session-specific set of 5 BCI trials that met this requirement from a repository of previous sessions. During a sham trial, we randomly selected a BCI trial from this fixed set and showed the visual feedback from that trial.

### Offline analysis of sham trials

To determine the efficacy of neurofeedback in counteracting fluctuations away from the activity target, we compared task performance in BCI trials with that of sham trials (Fig. 3 and Fig. 5). To assess the performance of sham trials, we passed the neural activity recorded during sham trials through the session’s BCI mapping in a post-hoc offline analysis. For each sham trial, we took bins during the entire 3.4s replay feedback period and applied the same decoding used during BCI feedback trials. The same procedure was applied to interleaved and consecutive sham trials.

Based on this offline analysis, we determined whether each sham trial was correct and computed success rates. We also applied a truncation scheme to subselect each sham trial’s bins for offline analysis (Fig. S2). We used only the retained bins to compute neural distances for sham trials. This truncation provided a more conservative test of BCI effectiveness. Including all time bins would have made sham performance appear worse, since untruncated sham trials had higher average neural distances than truncated sham trials (one-sided Wilcoxon paired signed rank test; *p <* 0.005 for each monkey; *N* = 21 sessions for each monkey).

### Computing average neural distances

To compare task performance in BCI and sham trials, we computed average neural distances at three levels: 1) Per-trial averages were computed by averaging across bins within each trial (Fig. 3 and Fig. 4). 2) Per-block averages were computed by averaging across bins within each trial and then across trials within each block (Fig. 3 and Fig. S3). 3) Per-condition averages were computed by averaging across bins within each trial, then across trials within each block, and finally across blocks within each session (Fig. 3, Fig. 5, and Fig. S4). For all three averaging procedures, only bins during the feedback period (i.e., excluding the fixation period) were included. For per-block and per-condition averages, both successful and unsuccessful trials were included, and only BCI trials were included for BCI blocks.

### Identifying changes in neural distance across trials

For each block of a session, we fit a linear regression model that captures the relationship between the clock time (the start time of each trial relative to the first trial in the block) and the per-trial average neural distance for that trial (Fig. 4A) :

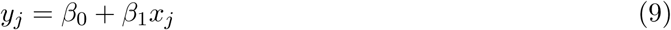

Here, *y_j_*∈ R represents the *j*-th trial’s per-trial average neural distance, *x_j_* ∈ R refers to the clock time for trial *j*, and *β*_0_*, β*_1_ ∈ R are parameters of the regression model.

Since BCI blocks lasted longer than sham blocks and natural fluctuations in neural activity away from the activity target accumulated over time, we matched block durations to avoid confounding slope differences between the two conditions. To achieve this, we end-aligned BCI blocks to their subsequent sham block and selected the last *k* BCI trials (both successful and unsuccessful) in the BCI block (interleaved sham trials were excluded) such that the clock time duration of these *k* trials matched the duration of the corresponding sham block (Fig. 4A). For instance, if a sham block lasted ∼ 90s, we selected the last *k* trials of the corresponding BCI block such that the *k* trials’ total time duration was roughly 90s. To account for long within-block pauses between trials that occurred when animals did not engage with the task, we separately computed the distribution of block clock-time durations for both BCI and sham blocks. We then excluded blocks with durations that were outliers relative to their respective condition’s distributions, defined as durations falling more than 1.5 times the interquartile range below the first quartile or above the third quartile, along with their paired blocks. For Monkey S, we removed 22 of 148 pairs of blocks (12 pairs due to BCI block outliers, 10 pairs due to sham block outliers). For Monkey P, we removed 33 of 211 pairs of blocks (16 due to BCI block outliers, 17 due to sham block outliers). Because BCI blocks contained more trials than sham blocks, animals had more opportunities to take breaks during BCI blocks, which may have contributed to the occurrence of BCI block-duration outliers. We also selected trials from the end of the BCI block, as they were the closest in time to the sham block trials that they would be compared to. For each session and condition (BCI block or sham block), we averaged the regression slope across all blocks of that condition (Fig. 4B).

### Comparing annulus diameter and pupil diameter

To assess whether animals regulated an arousal-related internal state during BCI usage, we quantified how closely changes in pupil diameter tracked changes in annulus diameter (Fig. 6). To compare pupil diameter (measured every 1 ms) to annulus diameter (computed every 50ms), we downsampled the pupil diameter measurements by using the median pupil diameter across samples within the corresponding 50ms bin. For each session, we then selected all the bins belonging to the feedback period on BCI trials (both successful and unsuccessful). We computed the Pearson correlation coefficient between the downsampled pupil diameters and annulus diameters.

For the shuffled distribution (Fig. 6), we computed the Pearson correlation between one session’s annulus diameters and another session’s pupil diameters. We considered every possible pair of the 42 sessions, yielding 861 (“42 choose 2”) correlation values. Since different sessions had varying numbers of BCI trials, we truncated the longer sessions to match the number of trials in the shorter sessions. Within each pairing of trials (across sessions), we truncated the longer trial, discarding any time points within the trial that occurred after the shorter paired trial ended.

## 1 Supplementary Figures

**Figure S1:**
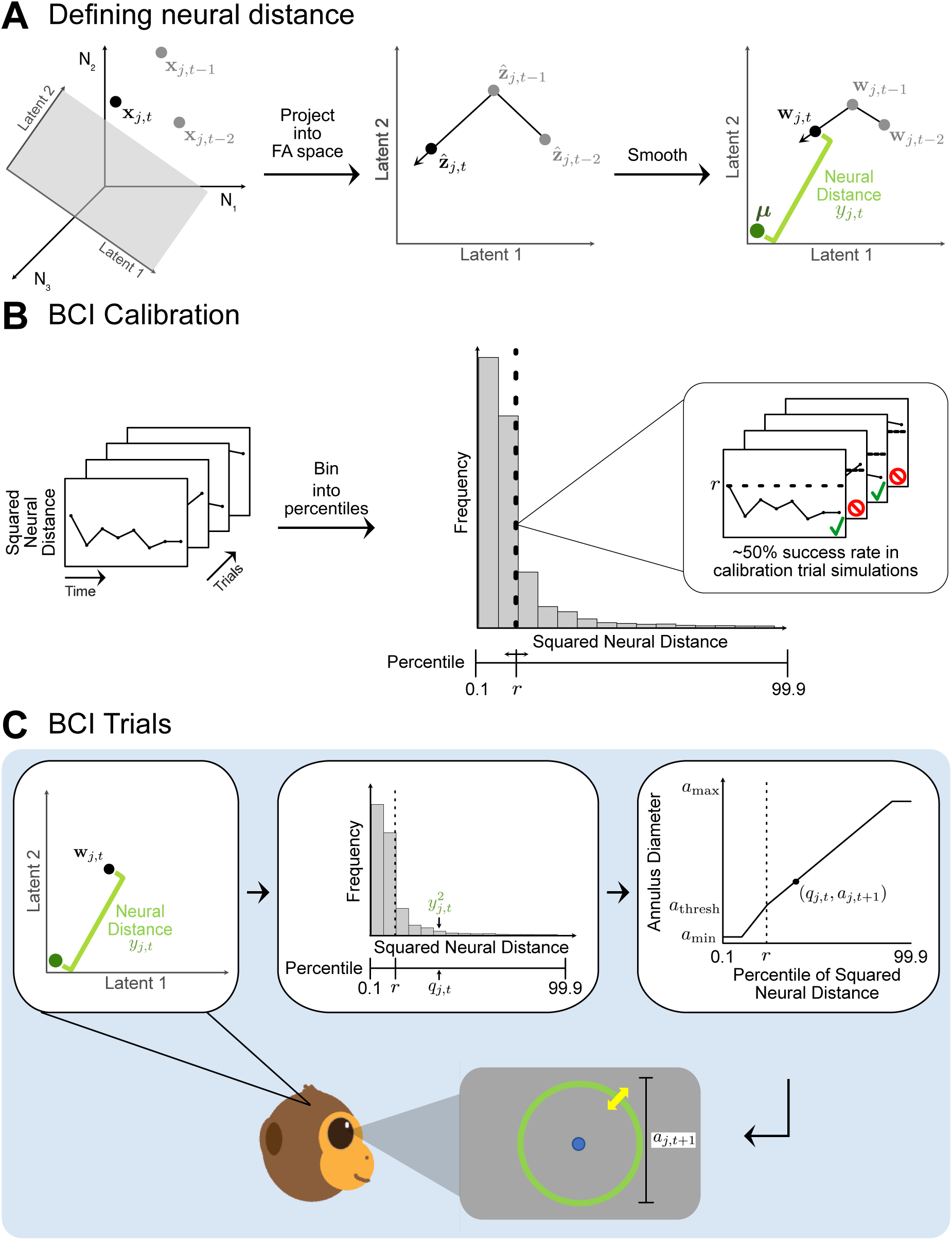
In BCI trials, the BCI mapping relates neural distance to visual feedback. **A.** Defining neural distance. To quantify how far the current neural activity is from an activity target, we first projected the spike count vector, **x***_j,t_* (black dot, left panel), into the FA subspace (gray plane, left panel) defined using the calibration trials (see Eq. 3). This yielded the FA projection for bin *t* of BCI trial *j*, **ẑ***_j,t_* (black dot, center panel). For illustrative purposes, we depict the FA subspace as a 2-dimensional plane, but in the actual experiments, we used higher-dimensional FA subspaces (see Methods). Second, we computed the smoothed FA projection, **w***_j,t_* (black dot, right panel), by applying exponential smoothing across time bins (Eq. 4). Lastly, we computed the neural distance, *y_j,t_* (green bracket, right panel), between the smoothed FA projection, **w***_j,t_*, and the activity target, ***µ*** (dark green dot, right panel), defined using the calibration trials (see Eq. 5). Gray dots represent the spike counts (left panel), FA projections (center panel), and smoothed FA projections (right panel) computed for previous time bins of the same trial. **B.** For the BCI calibration, we identified a reward threshold, *r*, that the animals needed to keep their neural distance below for 8 consecutive time bins (see Methods). We first computed the squared neural distances, *y*^2^, for bin *t* of calibration trial *i* across all bins of 60 calibration trials (stacked boxes, left panel; see Eq. 7). Second, we computed percentiles of the squared neural distances from 0.1 to 99.9 in increments of 0.1 (gray histogram, right panel). Lastly, we swept through the percentiles from 1 to 99 in steps of 1 and selected *r* (dashed line, right panel) to be the percentile increment corresponding to approximately 50% of the calibration trials being correct if they were processed as BCI trials (stacked boxes in the inset, right panel). We then used *r* as the reward threshold in the BCI mapping for all BCI trials in this session. **C.** During BCI trials, the BCI mapping determined the visual feedback based on the neural distance. For bin *t* of BCI trial *j*, we began by computing the neural distance, *y_j,t_*(green bracket, top left panel), using the smoothed FA projection, **w***_j,t_* (black dot, top left panel). We then identified the calibration-based percentile, *q_j,t_* (black arrow pointing to percentile line, top center panel), closest to the observed squared neural distance. Lastly, using a mapping that related *q_j,t_* to the annulus diameter (top right panel, see Eq. 8), we determined the annulus diameter for the next bin, *a_j,t_*_+1_ (black bracket, bottom gray panel).

**Figure S2:**
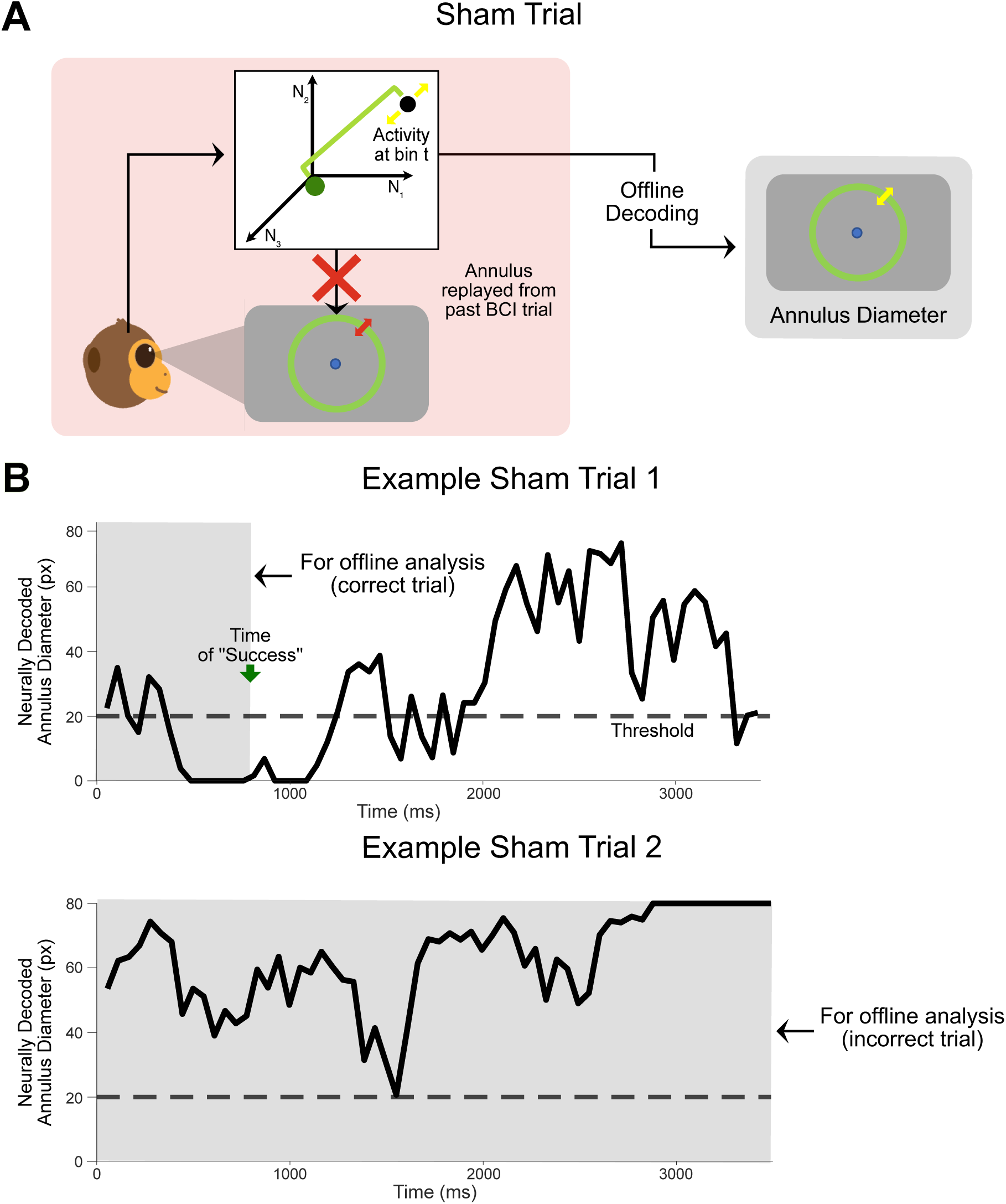
For all offline analyses, sham trials were processed like BCI trials. **A.** To compare a session’s BCI and Sham trials (Figs. 3 and 5), we passed neural activity recorded during sham trials (red box) through that session’s BCI mapping in a post-hoc offline analysis. For each sham trial (interleaved or consecutive sham), we used the neural activity recorded during the replay feedback period to decode the annulus diameters (see Methods). **B.** Truncation scheme. We asked if, at any point during the replay feedback period (0 – 3400 ms, divided into 68 bins of 50 ms), the annulus diameters decoded from neural activity recorded during this period met the success conditions for the BCI task. If so, we truncated the trial such that only neural activity from bin 1 to bin *t*, the earliest bin at which this success criterion was met (green arrow, example trial in top panel), was considered for offline analysis. This processing was equivalent to that for neural activity in BCI trials. Sham trials were only truncated if they were considered “correct” by the BCI mapping. If a sham trial was considered incorrect, we retained all the bins from that trial for further analysis (example trial in bottom panel). Shaded regions indicate the bins that were used for offline analysis. The black dashed line indicates the annulus diameter threshold (*a*_thresh_, see Methods) that consecutive bins had to remain below for success in the BCI task.

**Figure S3:**
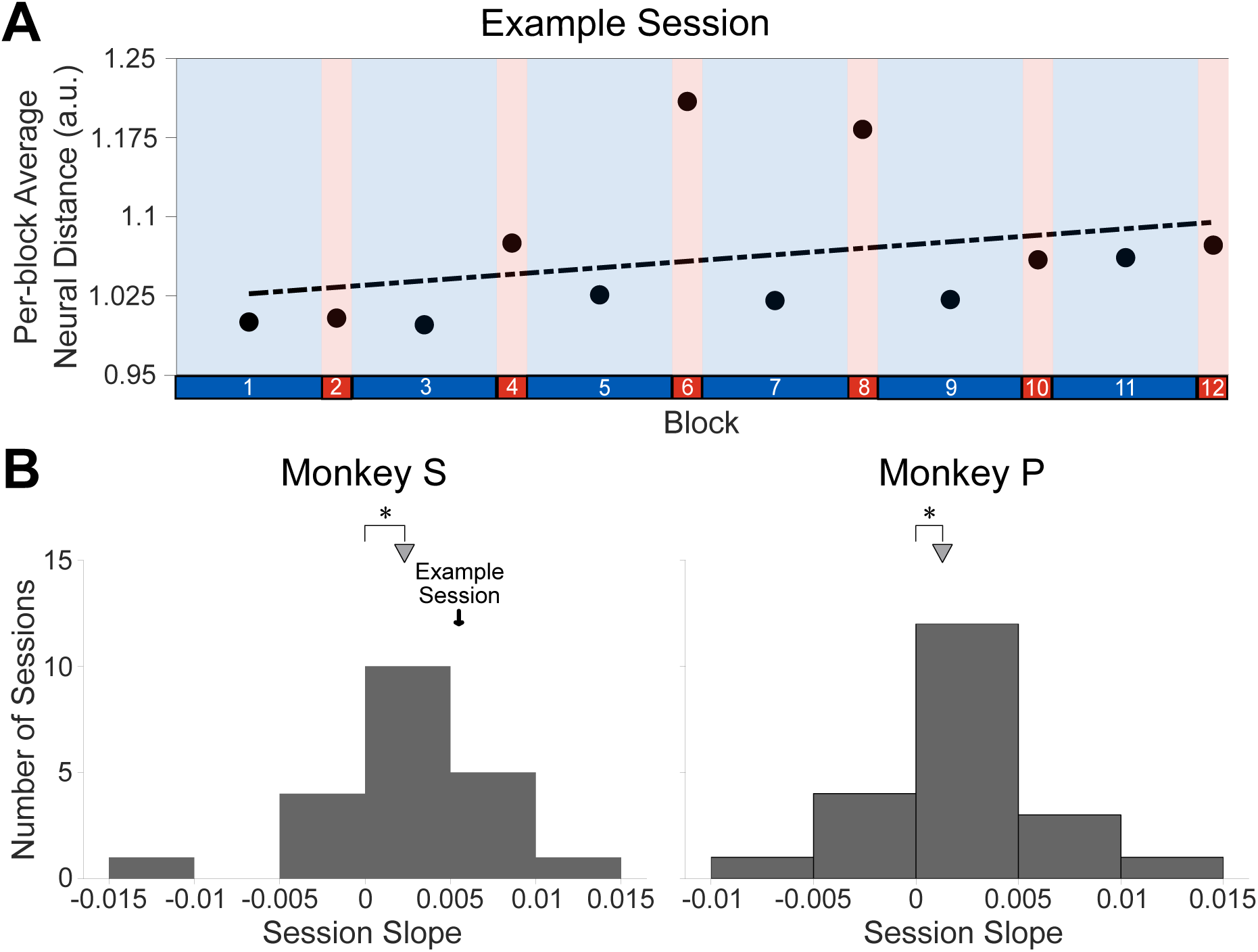
The BCI did not fully negate neural distance drifts across blocks within a session. Although Fig. 4 showed that BCI blocks counteracted across-trial fluctuations away from the activity target, we wondered whether the BCI blocks nullified the gradual distance increases that occurred within sham blocks. To investigate, we examined how the per-block average neural distance changed across blocks within a session. For each session, we fit a linear regression model relating the per-block average neural distance to block number, including both BCI and sham blocks in the regression. We found that within a session, neural distance still increased across blocks, indicating that BCI blocks did not fully rein in the distance drifts that occurred in sham blocks. **A.** In this example session (same as in Figs. 3 and 4), we saw an increase in the average neural distance across blocks (slope=0.0062). Interleaved sham trials were not included when computing the per-block average neural distance calculation for BCI blocks. The dashed line is the regression line between the per-block average neural distance (black dots, identical to horizontal lines plotted in Fig. 3B) and the block number within the session. **B.** The regression slopes were significantly positive for both animals (median of 0.0017 for sessions pooled from monkey S and monkey P; median of 0.0023 for Monkey S and 0.0013 for Monkey P; one-sided Wilcoxon signed rank test; *p <* 10*^−^*^3^ pooled across monkeys, *N* = 42 sessions; *p <* 0.05 for each monkey, *N* = 21 sessions for each monkey). Triangles indicate the median slopes observed across sessions.

**Figure S4:**
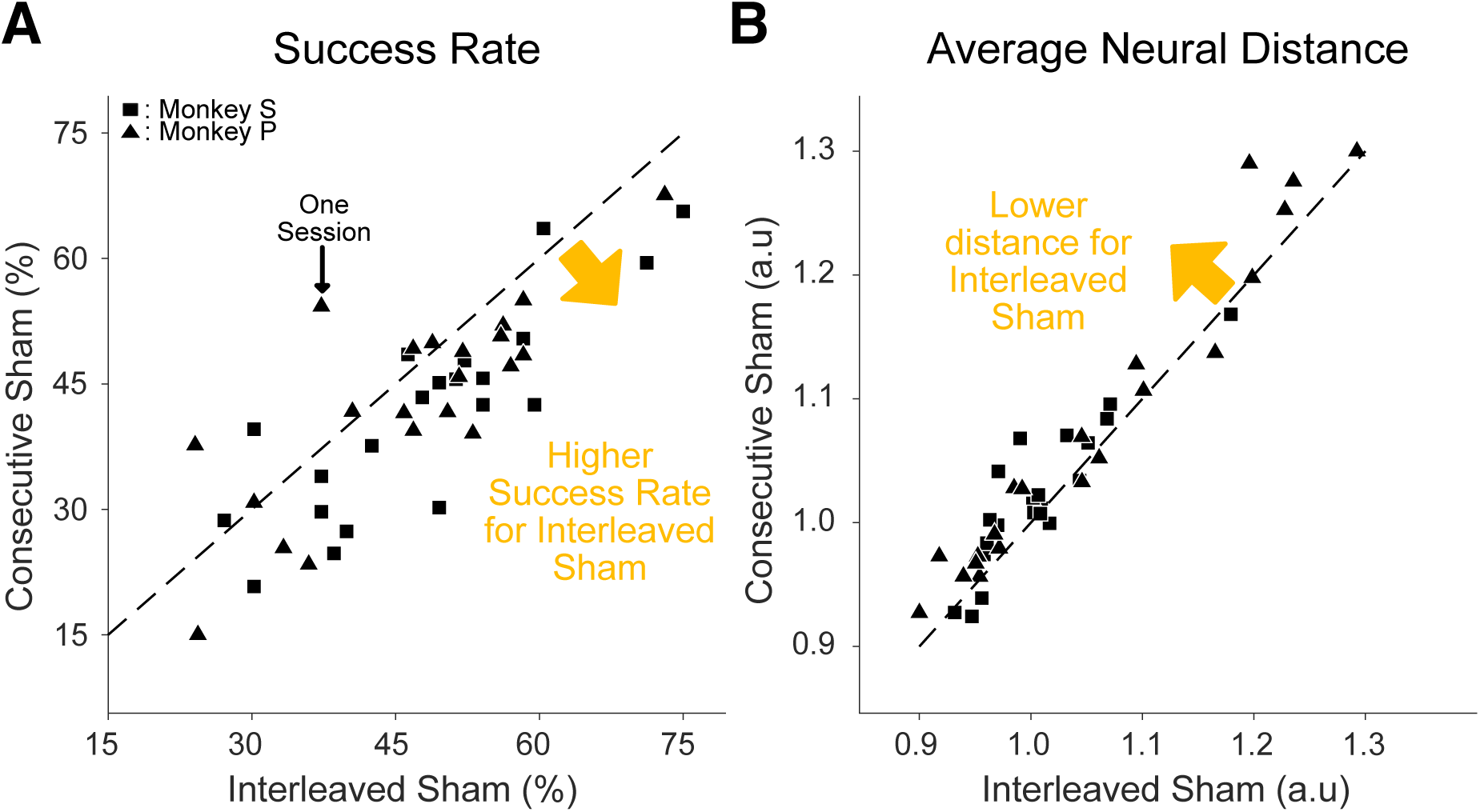
Subjects performed better in interleaved sham trials than in consecutive sham trials. In Fig. 5, we compared interleaved sham trials with BCI trials because, unlike consecutive sham trials, interleaved sham trials occurred within the same block context as BCI trials. Here, we tested whether trial context affected BCI performance and neural activity by comparing consecutive sham and interleaved sham trials. We found that animals performed better in interleaved sham trials than in consecutive sham trials. During interleaved sham trials, animals may have stayed engaged because these trials were interleaved with BCI trials. But during consecutive sham trials, they may have become disengaged once they realized their neural activity no longer controlled the annulus diameter. **A.** Interleaved sham trials had higher success rates than consecutive sham trials (one-sided Wilcoxon paired signed rank test; *p <* 10*^−^*^4^ pooled across monkeys, *N* = 42 sessions; *p <* 0.05 for each monkey; *N* = 21 sessions for each monkey). Success rates were computed by processing them like BCI trials (see Methods and Fig. S2). **B.** Per-condition average neural distances were lower for interleaved sham trials than consecutive sham trials (one-sided Wilcoxon paired signed rank test; *p <* 10*^−^*^4^ pooled across monkeys, *N* = 42 sessions; *p <* 0.05 for each monkey, *N* = 21 sessions for each monkey). When computing per-condition average neural distances for sham trials, we only included bins remaining after the truncation process described in Fig. S2 (see Methods). Each symbol represents the per-condition average neural distance for a session. Together, these results indicate that trial context affected BCI performance.

**Figure S5:**
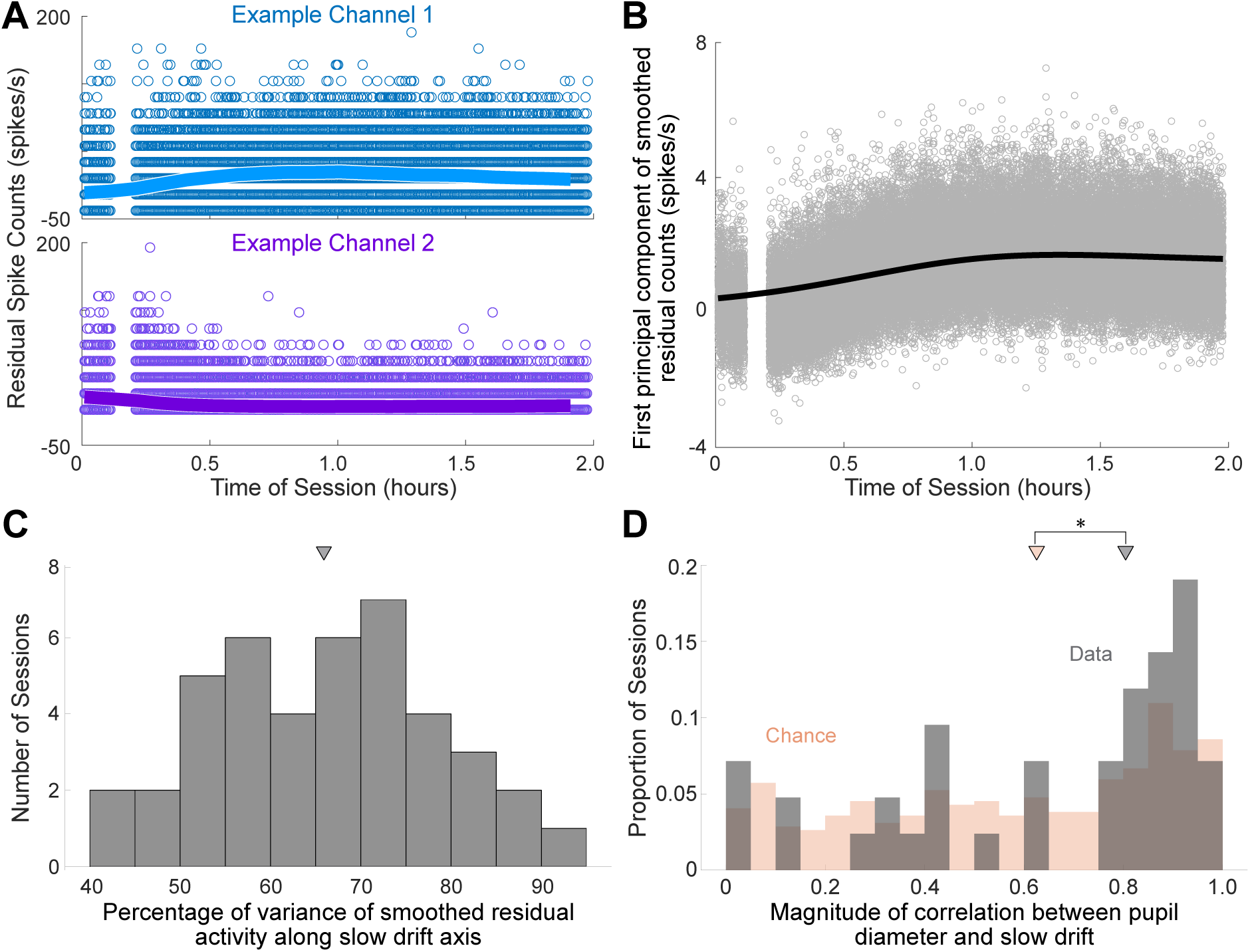
Slow changes in PFC activity are correlated with pupil-indexed changes in arousal. Here, we investigated whether changes in neural activity during BCI usage were correlated with arousal-related changes, as indicated by pupil diameter. **A.** Activity in individual channels fluctuated over time. Activity time courses for two example channels during a single session, showing residual counts from the fixation and feedback bins across successive trials that increase (top) or decrease (bottom) over time. Solid lines show residual spike counts boxcar-smoothed over 20-minute overlapping windows, with consecutive windows offset by 6 minutes, to highlight slow changes in activity. Each symbol represents a 50-ms-binned residual spike count, computed by subtracting each channel’s session mean from its raw spike count to remove differences in baseline firing rate across channels. **B.** Slow changes in population activity. From the same session in (A), this panel shows the time course of the linear combination of activity across 40 channels. This linear combination corresponds to a projection onto the “slow drift axis”, defined as the first principal component identified by applying PCA to residual spike counts boxcar-smoothed over 20-minute overlapping windows, with consecutive windows offset by 6 minutes, as described in (A). The parameters for the boxcar-smoothing, such as the window sizes and offsets, were chosen based on prior work that investigated slow changes in PFC neural activity (Cowley *et al.* [5]). Each gray symbol is the projection of a 50-ms-binned residual spike count onto this axis. For visualization only, the black solid line shows these projections after Gaussian smoothing with a 20-minute timescale. **C.** The slow drift axis captured most of the variance in slow population activity changes. The histogram shows the percentage of variance of boxcar-smoothed residual activity explained by the slow drift axis across sessions. For each session, we computed the % of variance of the boxcar-smoothed residual spike counts computed in **B** captured by the slow drift axis. We found that the slow drift axis explained a majority of slow activity changes per session (median of 65% for sessions pooled from both monkeys, represented by a triangle; medians of 61% for Monkey P and 71% for Monkey S). **D.** Slow changes in population activity are related to pupil diameter. For each session, we computed the running median of pupil diameter by taking the median within each 20-minute window, with consecutive windows offset by 6 minutes, to capture slow changes in pupil diameter. We also estimated the slow drift as the smoothed projections of the residual spike counts along the slow drift axis using the same window size and offsets. Then, we found the magnitude of the Pearson correlation coefficient over time between the running estimate of pupil diameter and slow drift for each session. We took the magnitude of correlations because PCA identifies sign-invariant axes (i.e., rotated up to 180 degrees), which makes the sign of slow drift arbitrary. The chance distribution was found by correlating the running medians of pupil diameter and smooth time courses randomly drawn from a Gaussian process. For the Gaussian process, we used the squared exponential covariance function with a timescale (45 mins) similar to that of slow drift, as reported in Cowley *et al.* [5]. We found that the correlations between pupil diameter and slow drift were larger (one-sided Wilcoxon rank-sum test; *p <* 0.05) when the slow drift was computed using neural recordings (gray histogram, median |*r*| = 0.8043) than when it was simulated (orange histogram, median |*r*| = 0.6265). This indicates that the observed correlations between pupil diameter and slow drift weren’t simply due to spurious correlations between two slow-varying signals. Triangles denote the medians of the distributions.

**Figure S6:**
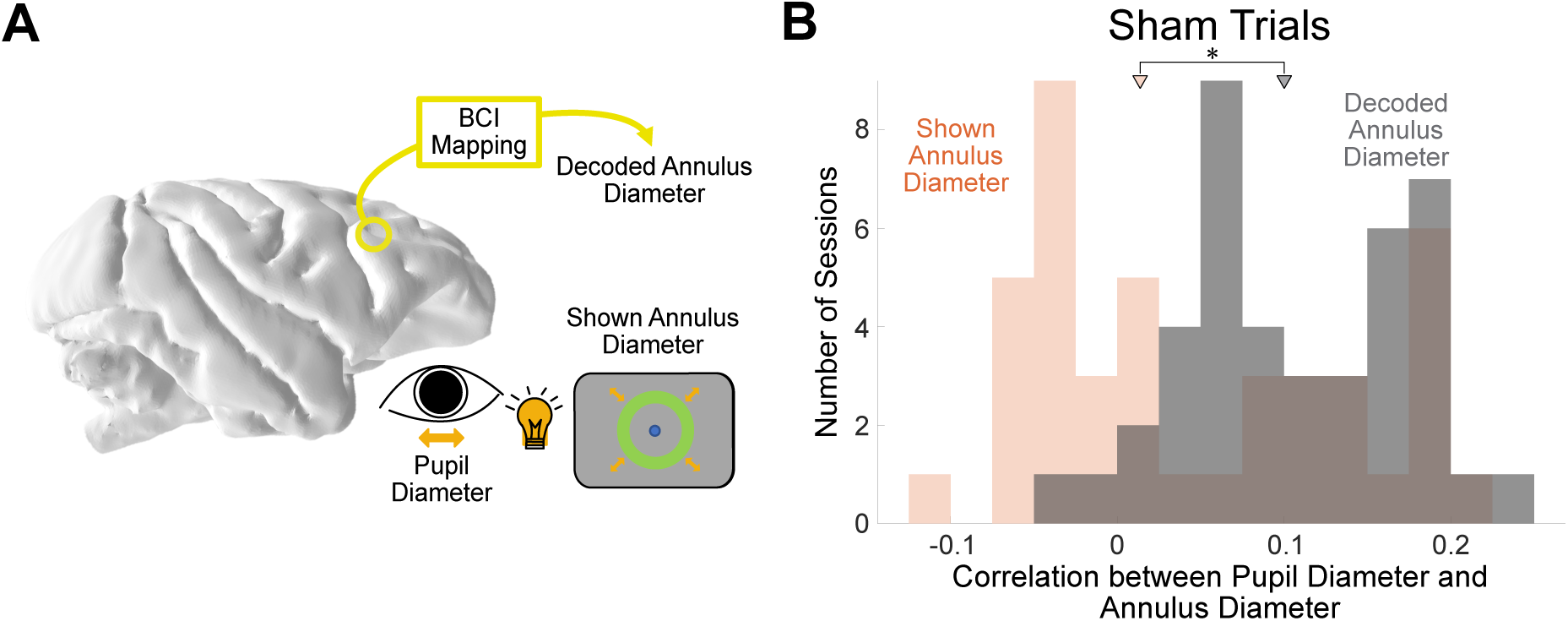
Luminance changes from the annulus did not solely drive pupil diameter changes. The positive correlation between pupil diameter and annulus diameter in BCI trials (Fig. 6B) might result from luminance changes rather than arousal-related changes. **A.** Annulus diameter changes may activate the pupillary light reflex: smaller annuli concentrate more luminance in the fovea, causing pupil constriction (Clarke *et al.* [71]). To test this hypothesis, we analyzed both interleaved and consecutive sham trials, which have two distinct annulus measures: decoded annulus diameter (obtained by passing neural activity recorded during replay feedback through the BCI mapping, Fig. S2) and shown annulus diameter (obtained from a trial of a previous session, see Fig. 2D and Methods). If the correlations observed in Fig. 6B were primarily due to luminance, pupil diameter should have more positive correlations with the shown annulus diameter than with the decoded annulus diameter in sham trials. This comparison cannot be made in BCI trials since the shown and decoded annulus diameters were identical. **B.** Luminance can not solely explain the changes in pupil diameters observed in Fig. 6B. For each session, we computed the Pearson correlation coefficient between the pupil diameter and the decoded annulus diameter for all bins during the feedback period (i.e., excluding the fixation period) in sham trials (median Pearson correlation coefficient *r* = 0.1005 for sessions pooled from both monkeys). For comparison, we also computed the Pearson correlation coefficient between the pupil diameter and the shown annulus diameter for all bins during the feedback period in sham trials (median *r* = 0.0136 for sessions pooled from both monkeys). For sham trials, the correlation was significantly higher (one-sided Wilcoxon rank sum test; *p* = 8.8406 × 10*^−^*^4^ pooled across monkeys, *N* = 42 sessions; *p* = 1.4351 × 10*^−^*^7^ for monkey S, *N* = 21 sessions; *p* = 0.1572 for monkey P, *N* = 21 sessions) for the decoded annulus diameter (gray histograms) than for the shown annulus diameter (orange histogram). Triangles refer to the median of each distribution.

## Acknowledgements

We are grateful to Samantha Nelson for help with animal training and data collection and to our animal care staff for their dedication in caring for the animals in this study. We are also grateful to the members of the Yu, Smith, Chase, and Batista laboratories for helpful discussions. The authors are inventors on a pending International Patent Application No. PCT/US2024/010801, which is related to the work presented in this paper. C.S.K was supported by the Richard King Mellon Presidential Fellowship and NIH T32 T32NS126122. B.M.Y. and M.A.S. were supported by NIH R01 MH118929, NSF NCS BCS 1734916/1954107, NSF NCS DRL 2124066/2123911, and NIH R01 EB026953.

